# Regulation of anterior neurectoderm specification and differentiation by BMP signaling in ascidians

**DOI:** 10.1101/2022.10.19.512847

**Authors:** Agnès Roure, Rafath Chowdhury, Sébastien Darras

## Abstract

Three palps make the most anterior structure of the ascidian larva. These ectodermal derivatives have both a sensory and adhesive functions essential for metamorphosis. They derive from the anterior neural border and their formation is regulated by signaling pathways such as FGF and Wnt. Since they also share gene expression profiles with vertebrate anterior neural tissue and cranial placodes, their study should shed light on the emergence of the unique vertebrate telencephalon. Here, we show that BMP signaling regulates two phases of palps formation in *Ciona intestinalis*. During gastrula stages, the anterior neural border marked by *Foxc* is specified in a domain of inactive BMP signaling, and activating BMP prevented its formation. Later on, inhibiting BMP led to the formation of a single large palp, most likely of dorsal identity. Our results indicate that BMP signaling regulates papilla *vs* inter-papilla fate decision within the palps forming region. Finally, we showed that modulating BMP signaling led to similar palps phenotypes in another ascidian species *Phallusia mammillata*. This led us to screen transcriptomic data and identify novel palps markers. Collectively, we provide a better molecular description of palps formation in ascidians that will be instrumental for comparative studies within ascidians and between ascidians and other chordates.

## Introduction

Ascidians (or sea squirts) belong to a group of marine invertebrates, the tunicates, that is the sister group of vertebrates. This phylogenetic position associated with a stereotyped embryonic development with few cells puts ascidians as interesting models for developmental biology and comparative approaches to address questions regarding chordates evolution and the emergence of vertebrates. Ascidians have a biphasic life cycle: following external development, the embryo gives rise to a swimming tadpole-like larva with typical chordate features (notochord, dorsal neural tube) that is going to attach to a substrate before metamorphosing into a sessile adult ascidian with a radically different body plan, a ‘bag’ with two siphons. Metamorphosis is controlled by a specific organ, the palps (also referred to as the adhesive organ or the adhesive papillae), that is located at the anterior end of the larva. The palps are a specialized part of the ectoderm that has adhesive and sensory properties. They enable the larva to select a suitable substrate for metamorphosis, hence a chemo- and/or mechano-sensory function, and to attach to it through the secretion of adhesive materials. It contains at least four cells types whose specification and function have not yet been deciphered in details (Johnson et al., 2020; Zeng et al., 2019). Three cell types - the ciliated sensory neurons, the collocytes (containing vesicles filled with adhesive material), and the axial columnar cells (ACCs) (myoepithelial cells controlling palps retraction following adhesion) - are elongated cells forming a protrusion. Three protrusions or papillae (two dorsal papillae that are bilaterally symmetrical, and a ventral papilla located at the midline; Fig 1C) make up the palps and are separated by the fourth cell type, the non-elongated inter-papillae cells.

**Figure 1.**
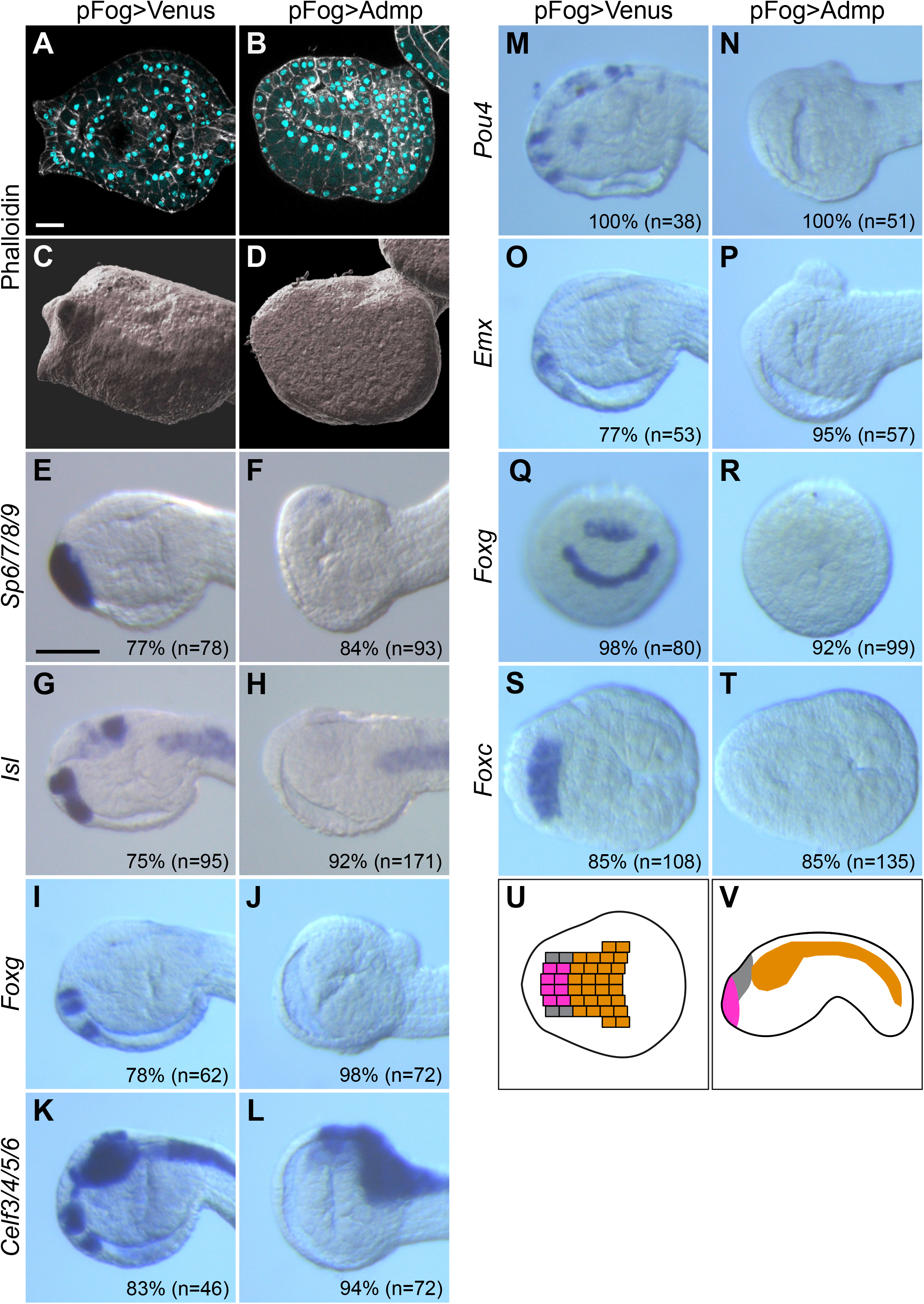
Early BMP activation prevents palps formation. BMP pathway was activated by overexpressing the BMP ligand Admp using the *Fog* ectodermal promoter. Experimental embryos were compared to control (overexpressing the fluorescent protein Venus). (A-D) Papilla protrusions and elongated cells were absent following BMP activation as revealed by confocal stacks for phalloidin (white) and DAPI (cyan) staining at larval stages (in A and B: confocal sections; in C and D: surface rendering). Scale bar: 20 μm. (E-T) BMP activation repressed genes expressed in the palps as determined by *in situ* hybridization for *Sp6/7/8/9* (E, F), *Isl* (G, H), *Foxg* (I, J), *Celf3/4/5/6* (K, L), *Pou4* (M, N) and *Emx* (O, P) at mid-tailbud stages (St. 23); and *Foxg* (Q, R) and *Foxc* (S, T) at neurula stages (St. 14). For each panel, n indicates the number of embryos examined. The percentages correspond to normal expression for pFog>Venus, and to gene repression in the palps territory for pFog>Admp. Experiments have been performed at least twice, except for *Celf3/4/5/6, Pou4* and *Emx* where results come from a single experiment. In tailbud embryos, a bulging mass of cells was often visible in the dorsal posterior trunk. It most likely corresponds to the CNS as revealed by *Celf3/4/5/6* expression that was outside of the embryo due to abnormal neural tube closure. Anterior to the left and dorsal to the top, except Q and R that are frontal views, and S and T that are neural plate views. Scale bar: 50 μm. (S, T) Schematic representation of the progeny of the neural plate: central nervous system in orange, palps region in magenta and aATENS precursors in gray.

Palps belong to the peripheral nervous system and have been instrumental for proposing evolutionary scenarios on the nervous system in chordates. In the ascidian *Ciona intestinalis*, palps cell lineage and topology, together with gene expression data and functional studies, have shown affinities with anterior derivatives of the vertebrate nervous system, the olfactory placodes and the telencephalon (Cao et al., 2019; Horie et al., 2018; Hudson et al., 2003; Liu and Satou, 2019; Poncelet and Shimeld, 2020; Thawani and Groves, 2020; Wagner and Levine, 2012; Wagner et al., 2014). Palps originate from precursors that are located at the anterior edge of the neural plate during gastrulation, that we will refer to the anterior neural border (ANB) (Fig 1U and 1V). While the ANB is not part of the central nervous system (CNS), it originates from the same lineage specified by FGF-mediated neural induction at the 32-cell stage and expresses neural markers such as *Celf3/4/5/6* (also known as *Etr* and *Celf3*.*a*) and *Otx* but does not express the epidermis factor *Tfap2-r*.*b* (also known as *Ap2-like2*). The separation between these two lineages is regulated by FGF/Erk signaling at gastrula/neurula stages, FGF being active in the CNS precursors. FGF signaling thus regulates positively and negatively two separate phases of palps specification. The ANB also expresses *Dmrt* and *Foxc*, coding for transcription factors that are essential for palps formation. From neurulation and through differentiation, palps express genes such as *Dlx*.*c, Foxg, Isl* or *Sp6/7/8/9* (also known as *Zf220* and *Btd*) whose orthologs specify anterior neural territories in vertebrates. In particular, *Foxg* and *Isl* are essential for palps formation. The ANB thus shares similarities with vertebrate anterior cranial placodes; and the palps share similarities with derivatives of the vertebrate telencephalon such as the olfactory bulb. It has been proposed that co-option of ANB/palps gene network to the anterior CNS led to the emergence of the vertebrate telencephalon (Cao et al., 2019).

While knowledge on transcription factors functions and interactions in palps formation has been elucidated in some details (Horie et al., 2018; Liu and Satou, 2019; Wagner et al., 2014), the role of cell-cell communication is scarce except for the involvement of FGF/Erk pathway (Hudson et al., 2003; Wagner and Levine, 2012). For example, we have previously shown that inhibition of canonical Wnt pathway is essential for ANB specification (Feinberg et al., 2019), but description of a function for Wnt signaling during later steps of palps formation is still lacking. Also, in a distantly related ascidian species *Halocynthia roretzi*, morphological data indicate that activating BMP pathway abolishes palps formation while BMP inhibition results in palps made of a single protrusion instead of three (Darras and Nishida, 2001). But the lack of molecular analysis prevents from precisely determining the function of BMP signaling.

We have directly addressed the function of BMP signaling pathway in palps formation during the embryogenesis of the ascidian *C. intestinalis*. We show that BMP is involved in two consecutive phases. Up to neurulation, ANB specification is incompatible with active BMP signaling; and the ANB forms in a region devoid of active BMP (as revealed by phospho-Smad1/5/8 immunostaining). Consequently, early activation of BMP prevents palps formation through the inhibition of ANB precursors formation. Following gastrulation, BMP participates in the differentiation of the palps through the regulation of the papillae *vs* inter-papillae fate decision. In particular, BMP-inhibited larvae harbor a single large protrusion made of elongated cells, the *Cyrano* phenotype, with an increased number of sensory neurons and ACCs. We propose that the competence to become a papilla is regulated by BMP through the transcription factors coding genes *Foxg* and *Sp6/7/8/9*. Interestingly, we show that modulating BMP pathway in the ascidian *Phallusia mammillata* (275 My of divergence time) produces the same phenotypes as in *C. intestinalis*. This allowed us to use previously published RNA-seq data (Chowdhury et al., 2022) to identify a number of novel genes expressed in the ANB and the palps. Altogether, our work points to a role for signaling pathways inhibition in ANB specification, similarly to early anterior neurectoderm formation in vertebrates. Moreover, we provide significant enrichment in palps gene network deciphering in order to probe its conservation with cranial placodes and telencephalon formation in vertebrates.

## Results

### BMP activation abolishes palps formation

When we activated the BMP signaling pathway by overexpressing, in the ectoderm, the BMP ligand Admp by electroporation using the pFog driver (Pasini et al., 2006), we observed an absence of protrusions that are an obvious features of the adhesive palps. The anterior end of the larvae were smooth, and epidermal cells were flat and did not display the typical elongated shape (Fig 1A-D). This morphological evidence was accompanied by the repression of the expression at mid-tailbud stages of all the genes expressed in the palps that we have examined: *Sp6/7/8/9, Isl, Foxg, Celf3/4/5/6, Pou4* and *Emx* (Fig 1 E-P). Palps derive from the median anterior neural border (ANB) at gastrula stages (Fig 1U and 1V) and can be tracked by the expression of genes essential for palps formation, *Foxg* at neurula stages and *Foxc* at gastrula stages (Fig 1 Q-T) (Liu and Satou, 2019; Wagner and Levine, 2012). Both genes were repressed by BMP activation; and this repression is sufficient to explain the later lack of palps gene expression and differentiation.

### Dynamic BMP activity in the palps forming region

The above results suggest that active BMP signaling is incompatible with palps formation. Active BMP signaling can be determined by examining the phosphorylated (active) form of the BMP transducer Smad1/5/8. It has been previously shown that BMP is active from late gastrula to early tailbud stages in the ventral epidermis midline of *C. robusta* embryos (Waki et al., 2015). We obtained similar results in *C. intestinalis* using a different antibody (Fig 2). More specifically, up to gastrulation, we did not detect significant levels for P-Smad1/5/8 except in a few posterior endomesodermal cells (Fig 2A). At mid-gastrula, P-Smad1/5/8 was present in the nuclei of the posterior (b-line) ventral midline epidermis (Fig 2B).

**Figure 2.**
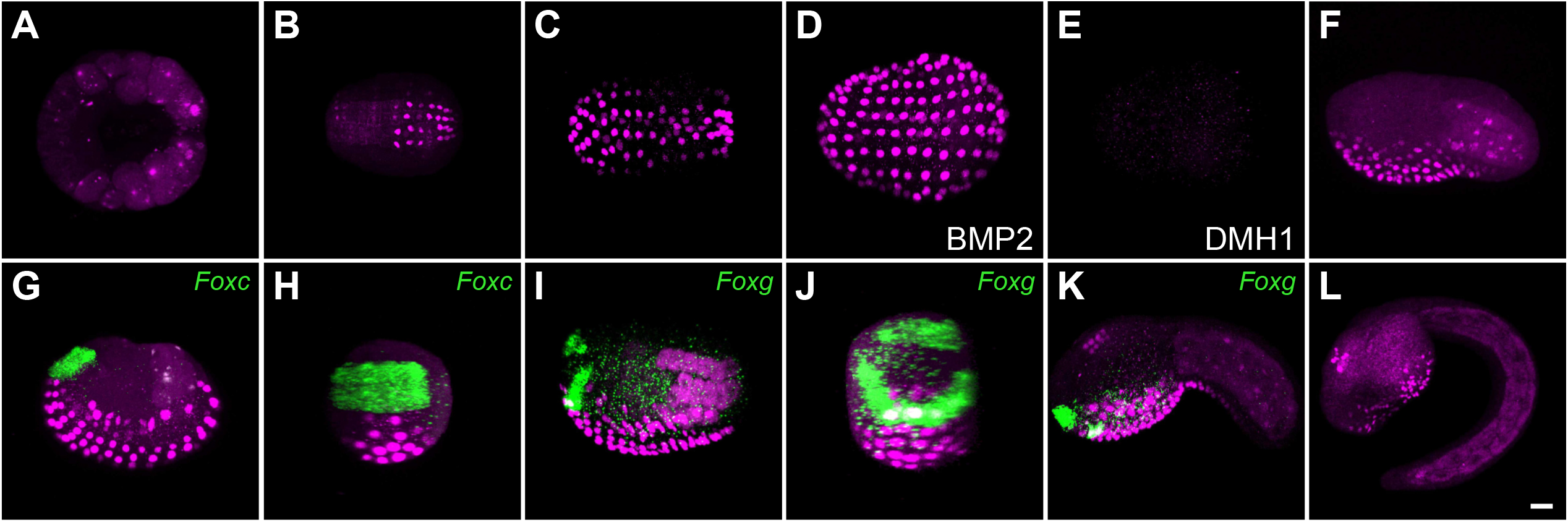
Dynamic BMP activity in the palps forming region. P-Smad1/5/8 immunostaining (magenta) was performed at various developmental stages: (A) early gastrula (St. 11), (B) late gastrula (St. 13), (C-E, G and H) early neurula (St. 14), (I, J) late neurula (St. 16), (F) initial tailbud (St. 18), (K) mid tailbud (St. 21) and (L) late tailbud (St. 23) stages. Gene expression by *in situ* hybridization was detected in green for *Foxc* (G and H) and *Foxg* (I-K). All embryos are control untreated except in D (100 ng/ml BMP2 protein treatment from the 8-cell stage) and E (2.5 μM DMH1 treatment from the 8-cell stage). Embryos are shown with anterior to the left with the following orientations: (A) vegetal view, (B-E) ventral views, (F, G, I, K and L) lateral views, and (H and J) frontal views. G and H are different views of the same embryo. I and J are different views of another embryo. All data have been obtained from at least two independent experiments. Scale bar: 20 μm.

Consequently, at the onset of *Foxc* expression in palps precursors, BMP is not active in the palps forming region. Shortly later, at early neurula stages, P-Smad1/5/8 extended into the anterior (a-line) ventral midline epidermis (Fig 2C). During neurulation, the posterior limit of P-Smad1/5/8 gradually shifted anteriorly in agreement with the dynamic posterior to anterior expression of candidate target genes (Roure and Darras, 2016); by mid-tailbud stages, P-Smad1/5/8 was restricted to the trunk ventral epidermis (Fig 2F, 2K and 2L). In addition, active Smad1/5/8 was also detected in the endoderm underlying the ventral epidermis midline with a similar temporal dynamic (Fig 2G, 2I, 2K and 2L), and in a group of cells of the anterior sensory vesicle at mid-tailbud stages (Fig 2K and 2L). We validated the specificity of these results by modulating BMP pathway: when embryos were treated with BMP2 protein, P-Smad1/5/8 was ectopically detected in the entire epidermis at early neurula stages, while no staining was observed following inhibition using the pharmacological inhibitor DMH1 (Fig 2D and 2E). To precisely relate the location of active signaling in the ectoderm and palps precursors, we performed double staining: P-Smad1/5/8 and *in situ* hybridization for the early palp markers *Foxc* and *Foxg* (Fig 2G-K). At late gastrula stages, P-Smad1/5/8 abutted *Foxc* expression domain, confirming that palps precursors were specified in a BMP-negative domain (Fig 2G and 2H). Later, we observed P-Smad1/5/8 in the median *Foxc* expression domain at mid neurula stages (not shown) and in the median part of the U-shaped *Foxg* expression domain (Fig 2I and 2J), corresponding presumably to the future ventral palp. This was confirmed by co-expression of P-Smad1/5/8 and *Foxg* in ventral cells in late neurulae and early tailbuds. (Fig 2J and 2K).

### BMP inhibition participates in ANB definition

We next tested whether BMP inhibition was sufficient to induce an ANB fate. When BMP signaling was blocked either by overexpression of the secreted inhibitor Noggin or by treatment with the BMP receptor inhibitor DMH1, *Foxc* expression at late gastrula stages was unchanged (Fig S1). The fact that *Foxc* was not ectopically expressed following BMP inhibition could be explained by an incomplete BMP blockade. However, DMH1 treatment led to undetectable P-Smad1/5/8 levels (Fig 2E). Alternatively, it could be that the number of cells that are competent to become ANB in response to BMP inhibition could be restricted to the cells already expressing *Foxc. Foxc* expression and palps fate are regulated by FGF signaling following neural induction and cell fate segregation (Wagner and Levine, 2012). We thus aimed at increasing the number of cells competent to form ANB by early activation of FGF signaling using treatment with recombinant bFGF protein, and testing the effects of BMP pathway modulations in this context. As expected, bFGF treatment from the 8-cell stage neuralized the entire ectoderm as revealed by the ectopic expression of the neural markers *Otx* and *Celf3/4/5/6* and the downregulation of the epidermal marker *Tfap2-r*.*b* at late gastrula stages (Fig 3 and S1). *Foxc* behaved somewhat unexpectedly: it was either ectopically expressed or repressed (Fig 3D). The repression of *Foxc* might be explained by the fact that FGF/Erk is downregulated in the palps lineage during gastrulation (Wagner and Levine, 2012), hence our continuous treatment might inhibit *Foxc* expression. Nevertheless, when BMP pathway was inhibited in addition to FGF activation, *Foxc* was strongly expressed as a cup covering the anterior end of the embryo, including the ventral epidermis (Fig 3F).

**Figure 3.**
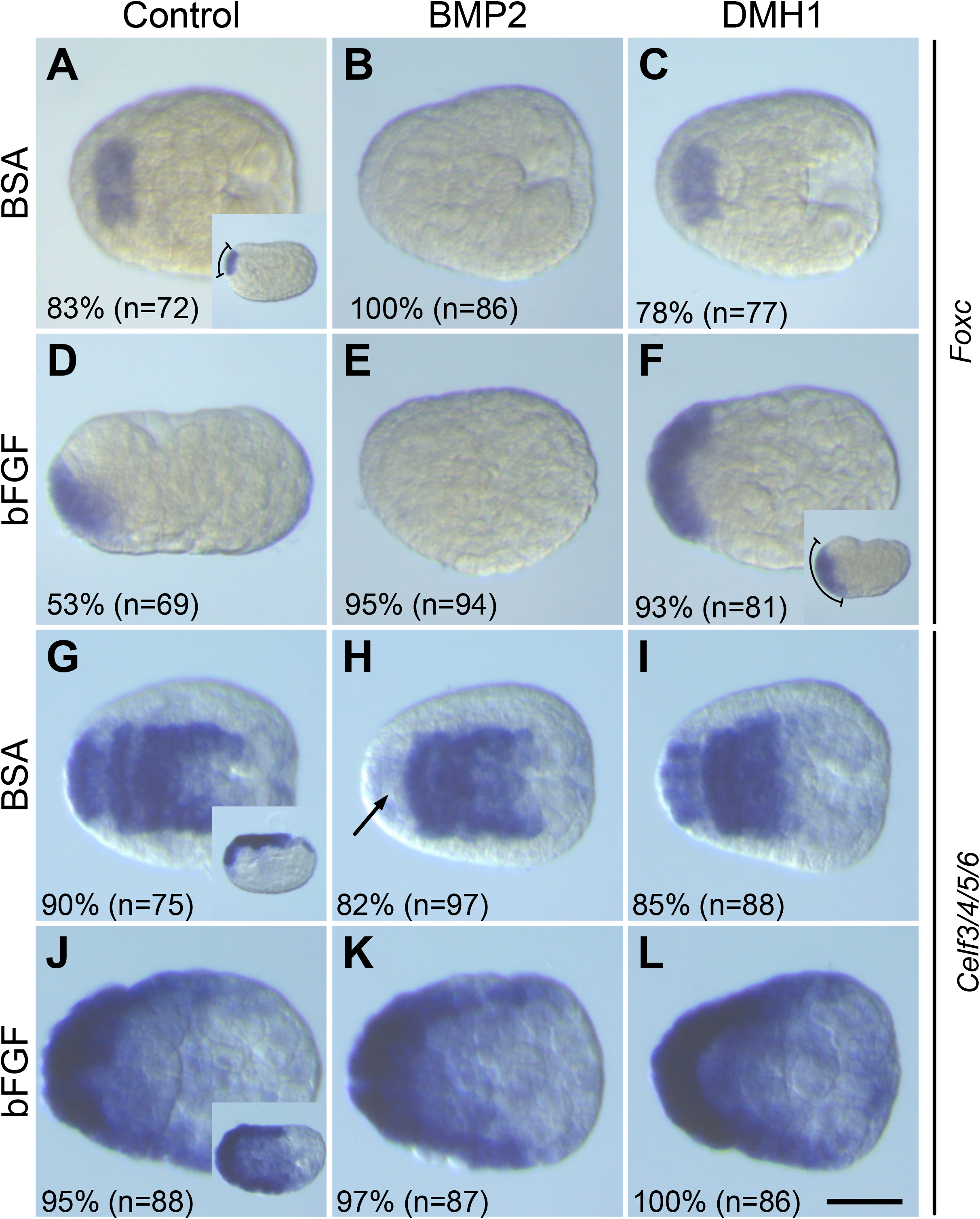
Inactive BMP signaling is required for ANB specification. Embryos were treated from the 8-cell stage to the fixation at early neurula stages (St. 14) with 100 ng/ml recombinant BMP2 protein or 2.5 μM DMH1 alone, or in combination with 100 ng/ml recombinant bFGF protein. Gene expression was assessed by *in situ* hybridization for *Foxc* (A-F) and *Celf3/4/5/6* (G-L). For each panel, n indicates the number of embryos examined. The percentages indicate the frequency of the phenotype depicted in the picture. The results come from two independent experiments. Embryos are shown in neural plate views with anterior to the left except insets that are lateral views with anterior to the left and dorsal to the top. Brackets in the insets in A and F highlight the dorso-ventral extension of *Foxc* expression. The arrow in H marks the downregulation of *Celf3/4/5/6* in the palps precursors. Scale bar: 50 μm.

Importantly, the loss of *Foxc* following BMP activation using recombinant BMP2 protein treatment was specific to this gene and did not result from neural tissue inhibition as in vertebrates since the neural markers *Otx* and *Celf3/4/5/6* were still expressed in the CNS but downregulated in the ANB (Fig 3 and S1). Reciprocally, BMP inhibition was not sufficient to lead to ectopic neural tissue formation. This is in agreement with similar data produced in the distantly related ascidian *Halocynthia roretzi* (Darras and Nishida, 2001). These observations show that BMP limits the domain of *Foxc* expression and ANB formation.

### The BMP signaling pathway regulates palps formation following ANB formation

We next examined whether modulating BMP had any impact on palps formation besides ANB specification. We thus performed whole embryo treatments starting at progressively later stages of embryonic development and examined the early marker *Foxc* and the late marker *Isl* (Fig 4). Activating BMP at early gastrula stages (St. 10) partially repressed *Foxc*, and abolished ventral palp formation. Mid-gastrula (St. 12) treatment had no effect on *Foxc* and repressed ventral palp at a lower frequency than the earlier treatment. Later treatments did not change *Isl* expression. Inhibiting BMP had no effect *Foxc* expression as for the earliest treatment (Fig 3 and S1). *Isl*, that is normally expressed as 3 spots, had a U-shaped expression. This phenotype was much less frequent when the DMH1 treatment started at late neurula stages (St. 16).

**Figure 4.**
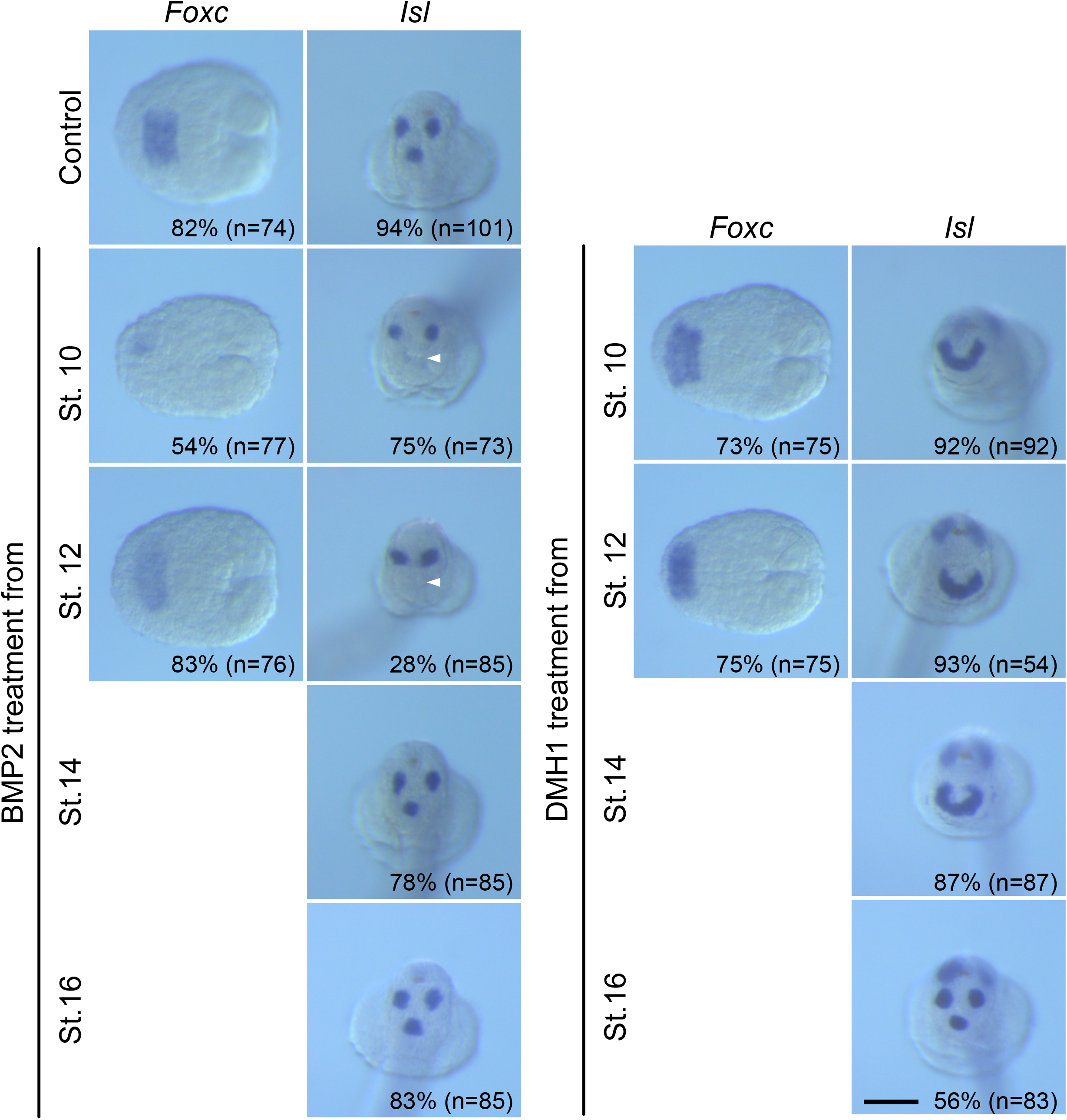
Late effects of BMP pathway modulations on palps formation. Embryos were treated with 100 ng/ml recombinant BMP2 protein (left panels) or 2.5 μM DMH1 (right panels) from the stage indicated on the figure up to fixation and *in situ* hybridization for *Foxc* at early neurula stages (St. 14) and *Isl* at late tailbud stages (St. 23). For each panel, n indicates the number of embryos examined. The percentages indicate the frequency of the phenotype depicted in the picture. The results come from two or more independent experiments. Embryos are shown in neural plate views with anterior to the left for *Foxc* and frontal view with dorsal to the top for *Isl*. White arrowheads highlight the absence of the ventral spot of *Isl* expression. Scale bar: 50 μm.

We next focused on understanding the U-shape pattern. It reminded us endogenous *Foxg* expression (Liu and Satou, 2019): at neurula stages, *Foxg* was expressed in the future palps following a U-shape that gradually converted into a 3-spots pattern at tailbud stages that prefigures the three papillae protrusions (Fig 5A-D). It thus seems that inhibiting BMP prevents the refinement of *Foxg* expression. Accordingly, *Foxg* expression was U-shaped following DMH1 treatment (Fig 5E). Interestingly, knockdown of *Sp6/7/8/9* leads to a U-shaped *Foxg* expression (Liu and Satou, 2019). Since *Sp6/7/8/9* and *Foxg* are initially partially co-expressed before showing exclusive patterns, it has been proposed that *Foxg* restriction to the future protrusions is the result of repression by Sp6/7/8/9. We thus determined *Sp6/7/8/9* and *Foxg* expression following BMP modulation from early gastrula stages (Fig 5). When embryos were treated with BMP2 protein, the ventral expression of *Foxg* in the U-shape was missing and *Sp6/7/8/9* was ectopically expressed at this location at late neurula/initial tailbud stages (Fig 5G and 5J). Reciprocally, following DMH1 treatment, ventral *Sp6/7/8/9* expression was shifted to a median position while *Foxg* was unchanged (Fig 5H and 5K). Consequently, the palps-forming U-shape most likely does not contain *Sp6/7/8/9* positive cells. In conclusion, we propose that the papilla *vs* inter-papilla fate choice is controlled by BMP signaling through the regulation of the expression of *Sp6/7/8/9*.

**Figure 5.**
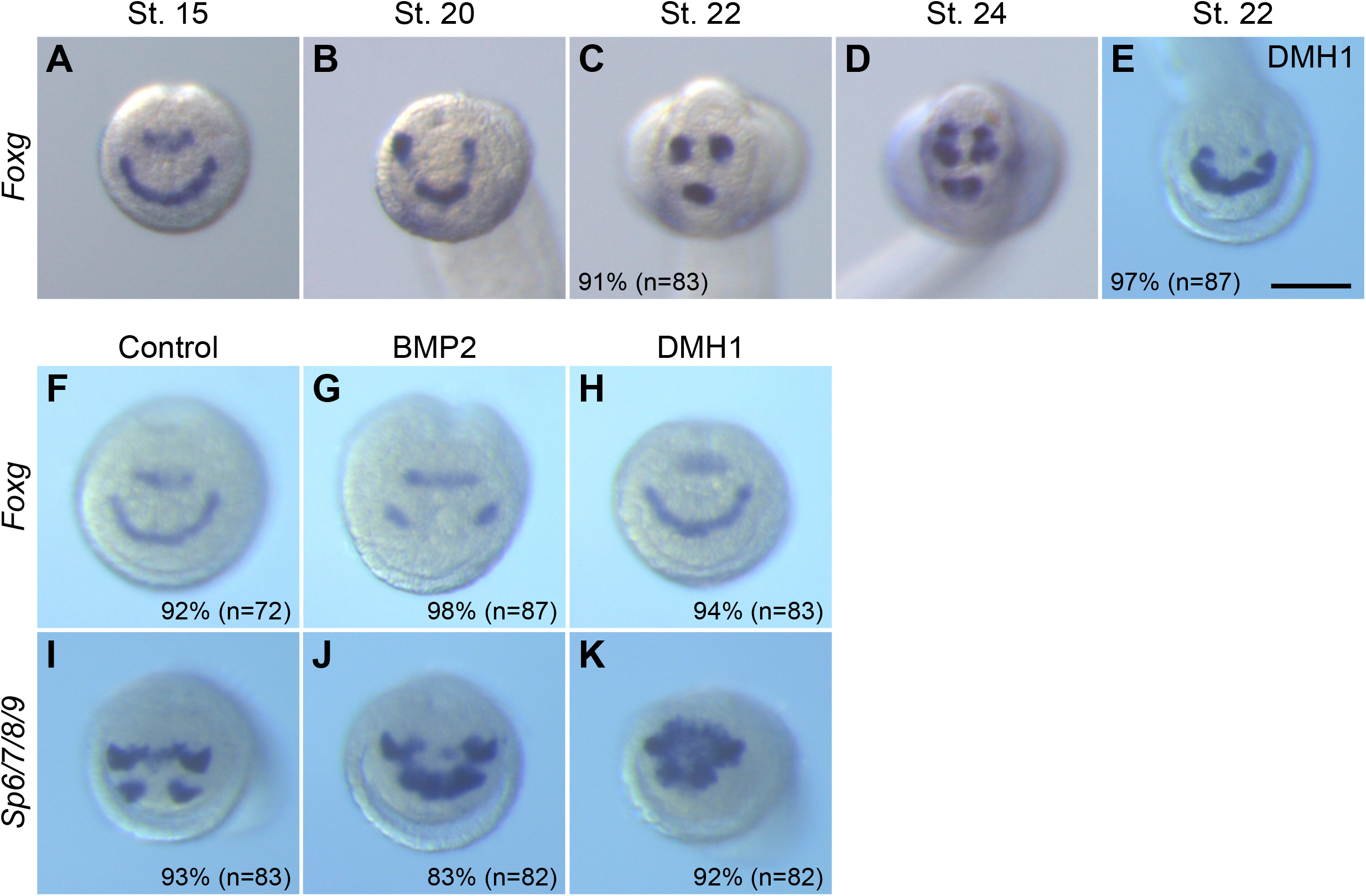
Regulation of *Foxg* and *Sp6/7/8/9* by BMP signaling. (A-D) Expression of *Foxg* at different developmental stages in the palps forming area (the specific stage is indicated at the top of each picture). Note that, at early stages (A), *Foxg* was expressed in two anterior ectodermal territories, the U-shaped palps forming region and a more dorsal row of cells likely contributing to the oral siphon primordium (Liu and Satou, 2019). (E) *Foxg* was expressed following a U-shape when BMP pathway was inhibited from early gastrula (St. 10) using 2.5 μM DMH1. (F-K) Expression of *Foxg* (F-H) and *Sp6/7/8/9* (I-K) in control embryos (F and I), embryos treated with 100 ng/ml BMP2 protein (G and J), and embryos treated with 2.5 μM DMH1 (H and K) from early gastrula (St. 10). For each panel, n indicates the number of embryos examined. The percentages indicate the frequency of the phenotype depicted in the picture. The results come from two independent experiments. Embryos are shown in frontal view with dorsal to the top. Scale bar: 50 μm.

### A single protrusion with additional neurons following BMP inhibition

We examined how the *Isl*/*Foxg* U-shape differentiated. While *Isl* was expressed following a large U at mid-tailbud stages (St. 23) (Fig 4), it was concentrated in a protruding structure at the anterior tip of late tailbuds (St. 25) (Fig 6A and 6F). In larvae, this single large protrusion was made of elongated cells as visualized by phalloidin staining (Fig 6B, 6C, 6G and 6H). We coined this phenotype *Cyrano* (in memory of the famous character depicted by Edmond Rostand). In *Ciona*, it is thought that palps contain a fixed number of the different cell types (Zeng et al., 2019). We performed fluorescent *in situ* hybridization at late-tailbud stages (St. 23) using four genes and made 3D reconstruction of the z-stacks acquired by confocal microscopy (see Material and Methods) in order to determine the differentiation of the palps in the *Cyrano* embryos. In agreement with previous reports, we found that the ACC marker *Isl* was expressed in 12 cells in control embryos (4 cells par papilla) (Fig 6D, 6E and 6K). By contrast, we found that the sensory neuron marker *Pou4* was expressed in 10 cells instead of 12 (Fig 6D, 6E and 6K). Interestingly, both dorsal palps had 4 cells that surrounded the *Isl*-positive cells while the ventral palp contained 2 *Pou4*-positive cells located dorsally to the *Isl*-positive cells. In DMH1 treated embryos, the number of neurons increased to 18 on average and the number of ACCs to 18 (Fig 6K). We also observed an increase of *Cel3/4/5/6* cells. Interestingly, the number of cells expressing *Sp6/7/8/9*, that has been described as an inter-papillae marker (Wagner et al., 2014), was decreased in DMH1 treated embryos (Fig 6K). These data are in agreement with the interpretation that the number of cells with papilla fate has increased, however their physical proximity likely leads to the formation of a single protrusion.

**Figure 6.**
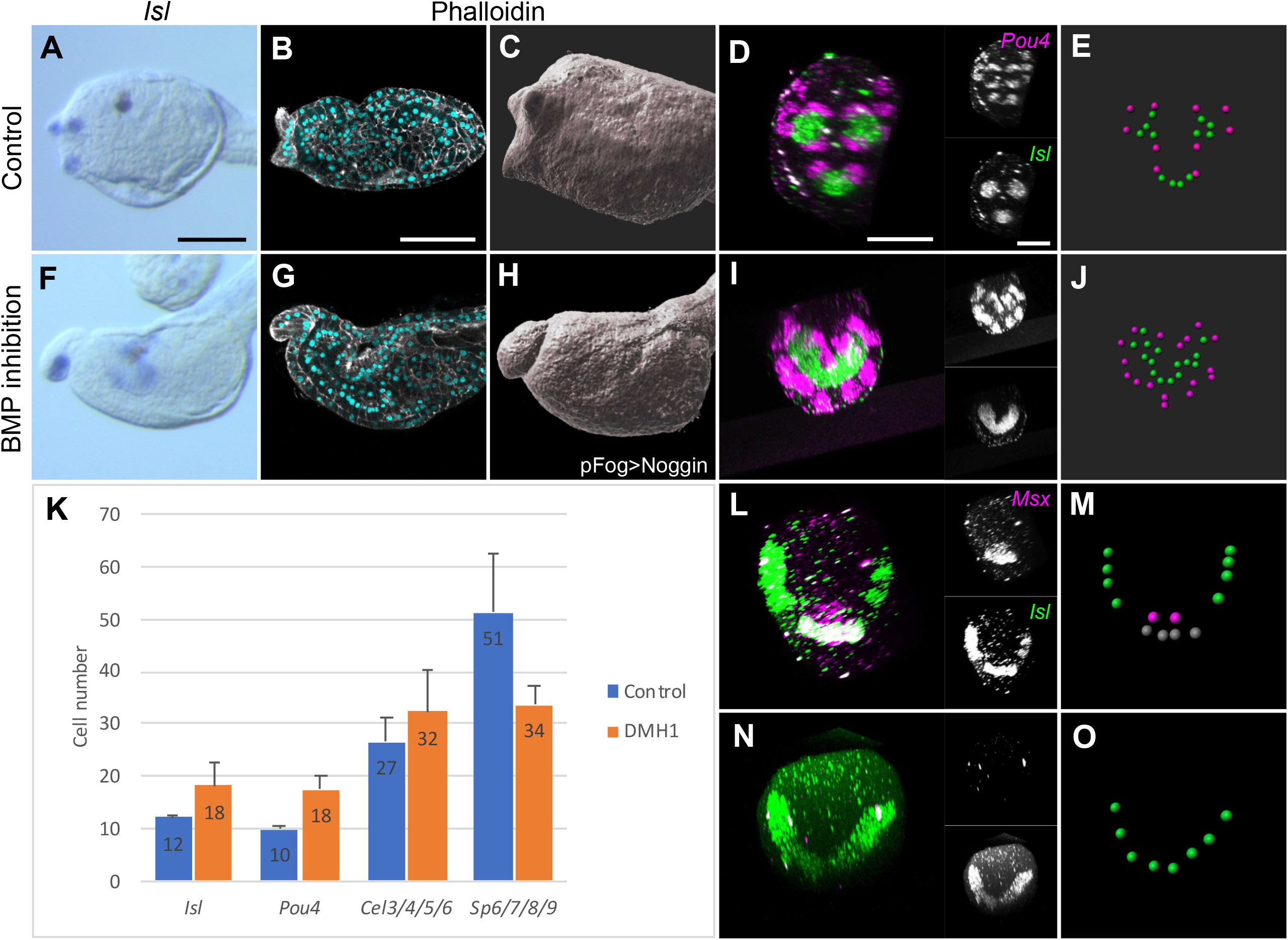
BMP inhibition leads to the formation of a single large palp of dorsal character. Embryos were BMP signaling was inhibited using treatment with 2.5 μM DMH1 from St. 10 (F, G, I, J, N and O) or Noggin overexpression (H) were compared to control embryos for morphology and gene expression by *in situ* hybridization. *Isl*, normally expressed in each of the 3 protruding palps (A, 86%, n=7), was expressed as a large spot in a single protrusion at late tailbud stages (St. 25) in treated embryos (F, 100%, n=21). The single large protrusion is made of elongated cells (B, C, G and H; DAPI in cyan and phalloidin in white). Double fluorescent *in situ* hybridization for *Pou4* (magenta) and *Isl* (green) in control (D) and treated embryo (I) at late tailbud stages (St. 23). 3D representation of nuclei for cells expressing each gene (E and J). (K) Count of the number of cells expressing each gene at late tailbud stages (St. 23) using 3D reconstructions as in E and J. The graph represents the average values from two or more independent experiments, with error bars denoting the standard deviation. The numbers of embryos examined are as follows: control embryos (*Isl*: 14, *Pou4*: 5, *Celf3/4/5/6*: 5, and *Sp6/7/8/9*: 6) and DMH1 treated embryos (*Isl*: 9, *Pou4*: 6, *Celf3/4/5/6*: 5, and *Sp6/7/8/9*: 5). Double fluorescent *in situ* hybridization for *Msx* (magenta) and *Isl* (green) in control (L) and treated embryo (N) at early tailbud stages (St. 19). 3D representation of nuclei for cells expressing each gene (M and O). Embryos are shown with dorsal to the top in lateral view (A-C and F-H) or frontal view (D-E, I-J, and L-O). Scale bars: 50 μm.

In DMH1 embryos, *Pou4* was expressed all around the *Isl* cells like in the dorsal palps. This suggested that the *Cyrano* protrusion may have a dorsal identity. In support of this interpretation, we found that the expression of the homeobox transcription factor *Msx*, that we found transiently expressed in the future ventral palp at the onset of *Isl* expression (Fig 6L and 6M), was lost following BMP inhibition (Fig 6N and 6O).

### Palps formation is similarly regulated by BMP in *P. mammillata*

We aimed at determining the conservation of the role of BMP in palps differentiation by examining embryos of the ascidian *P. mammillata* that belongs to the same family as *Ciona*, the Phlebobranchia, but with a significant divergence time (275 My) (Fig 7A) (Delsuc et al., 2018). First, we determined that BMP signaling was active is the ventral part of the embryo with a similar dynamic to *Ciona* as revealed by P-Smad1/5/8 immunostaining (Fig S2). Next, we identified single orthologs for *Celf3/4/5/6, Pou4* and *Isl* genes, that were all expressed in the palps (Fig 7) (Chowdhury et al., 2022; Coulcher et al., 2020; Dardaillon et al., 2020).

**Figure 7.**
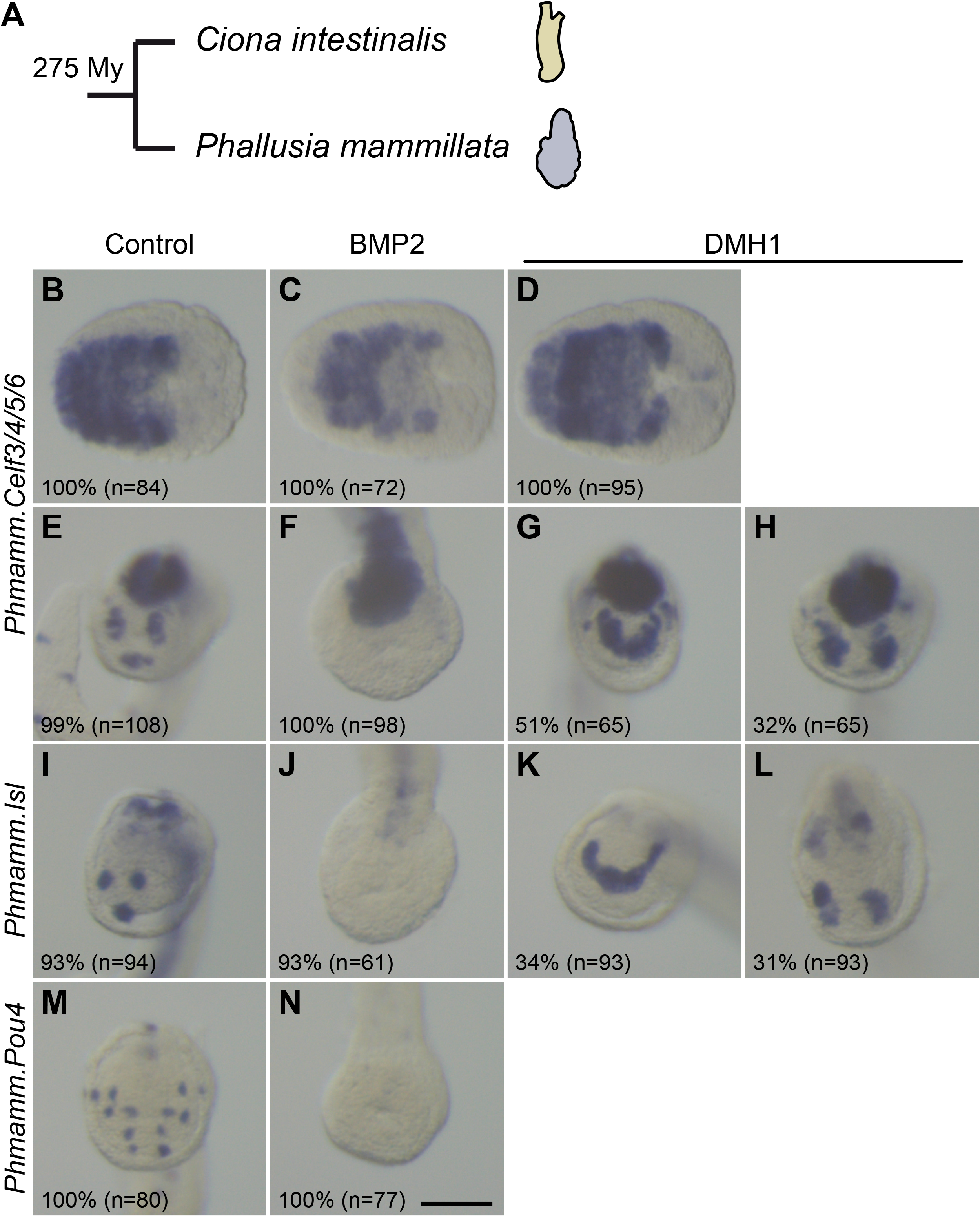
The BMP signaling pathway regulates palps formation in *Phallusia mammillata*. (A) Schematic representation of the appearance of adults *C. intestinalis* and *P. mammillata*, and their phylogenetic distance. (B-N) Modulating the BMP pathway modifies gene expression in the palps. *P. mammillata* embryos were treated from the 8-cell stage with 150 ng/ml recombinant BMP2 protein (C, F, J and N) or 2.5 μM DMH1 (D, G, H, K and L). They were fixed at neurula stages (B-D) and mid/late tailbud stages (E-N). Expression patterns for *Celf3/4/5/6* (B-H), *Isl* (I-L) and *Pou4* (M, N) was determined by *in situ* hybridization. For each panel, n indicates the number of embryos examined. The percentages indicate the frequency of the phenotype depicted in the picture. The results come from two or more independent experiments. Embryos are shown with anterior to the left in neural plate view (B-D), and in frontal view with dorsal to the top (E-N). Scale bar: 50 μm.

Treatment with recombinant BMP2 protein from the 8-cell stage abolished expression of all three markers in the palps, like in *Ciona* (Fig 7). Following DMH1 treatment from the 8-cell stage, both *Celf3/4/5/6* and *Isl* were expressed in the palps territory following a U-shape pattern like in *Ciona* but not in all cases. For a large fraction of embryos, the pattern appeared as two bars of intense staining resembling the U-shape but without the ventral part. This phenotype that we did not observe in *Ciona* might reveal some differences in the role of BMP in the two species.

Given the overall similar effects on palps formation after alterations of BMP signaling, we sought to identify novel palps molecular markers by using a dataset previously generated in *P. mammillata* (Chowdhury et al., 2022). We had generated, at several developmental stages, RNA-seq data for whole embryos treated with recombinant BMP4 protein and/or DAPT, a pharmacological Notch inhibitor. We identified 1098 genes repressed by BMP signaling at least at one developmental stage (Table S1). In this list, we found the orthologs for 11 well defined *Ciona* palps markers; and 4 of them (*Otx, Isl, Atoh1/7* and *Celf3/4/5/6*) were described as expressed in the palps lineage in *Phallusia* (Coulcher et al., 2020; Dardaillon et al., 2020). Using Gene Ontology analysis, we selected a list of 53 genes encoding developmental regulators (transcription factors and signaling molecules) or involved in neural tissue formation, and examined their expression patterns (Table S2). By searching the Aniseed database (Dardaillon et al., 2020) and our previously generated expression data (Chowdhury et al., 2022; Coulcher et al., 2020), we identified 12/26 genes expressed in the palps. By performing *in situ* hybridization for 27 extra genes, we discovered 7 novel palps markers whose expression is shown in Fig 8.

**Figure 8.**
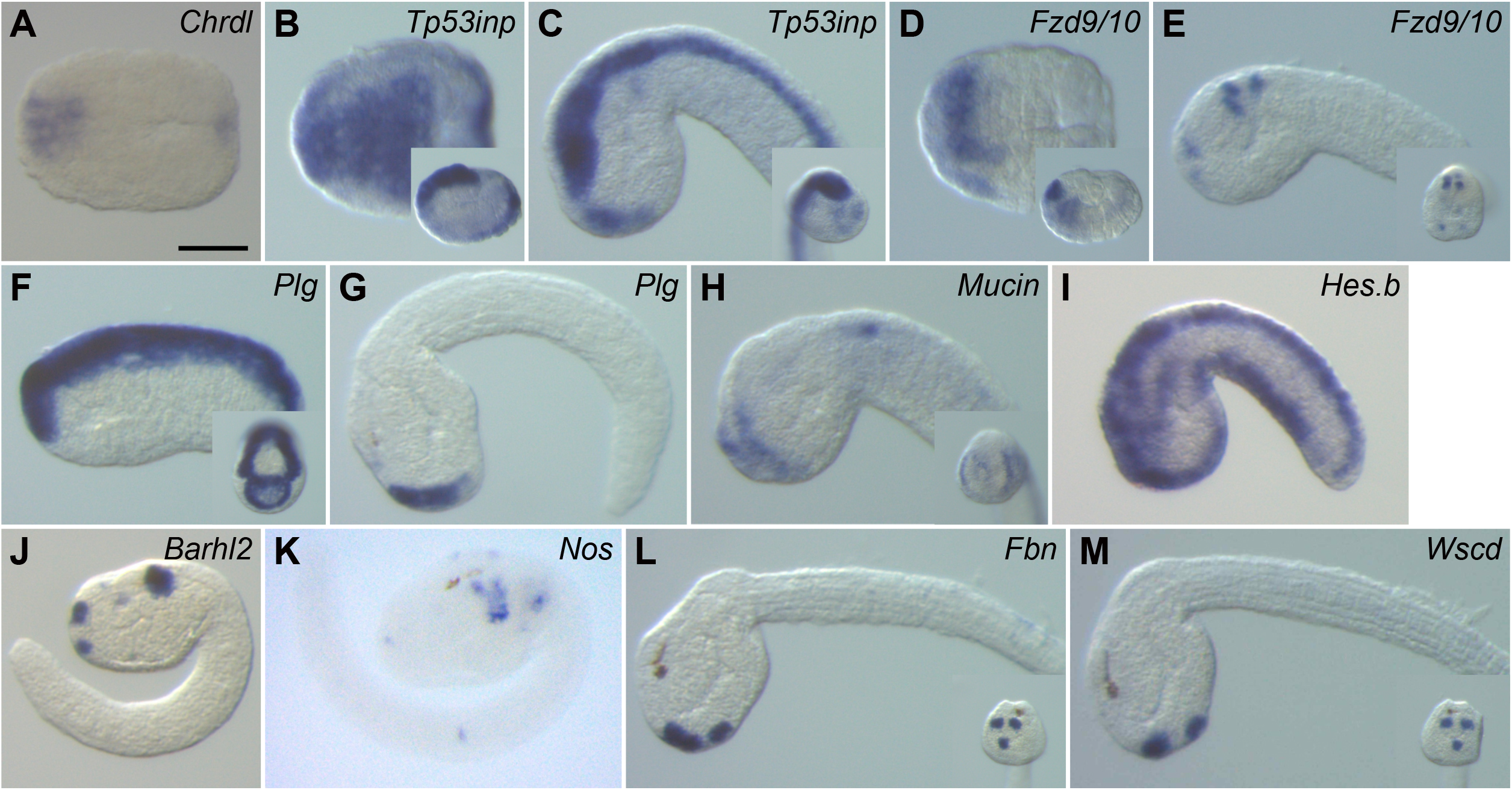
Identification of genes expressed in the palps in *Phallusia mammillata*. *In situ* hybridization at selected stages for *Chrdl* (A), *Tp53inp* (B, C), *Fzd9/10* (D, E), *Plg* (F, G), *Mucin* (H), *Hes*.*b* (I), *Barhl2* (J), *Nos* (K), *Fbn* (L) and *Wscd* (M). Embryos are shown with anterior to the left in neural plate view (A, B and D), in lateral view with dorsal to the top (B inset, C, D inset, E-M), and in frontal view with dorsal to the top (insets in C, E, F, H, L and M). Scale bar: 50 μm.

Surprisingly, by examining the expression data generated previously, we found that some genes with palps expression were up-regulated by BMP in our dataset, such as *Chrdl* and *Nos* (Table S2 and Fig 8). To have a broader view of the potential effect of BMP signaling on gene regulation in the palps, we gathered, from previous publications (Chen et al., 2011; Chowdhury et al., 2022; Coulcher et al., 2020; Joyce Tang et al., 2013; Kusakabe et al., 2012; Liu and Satou, 2019; Pasini et al., 2006; Roure and Darras, 2016; Shimeld et al., 2005; Wagner and Levine, 2012; Wagner et al., 2014), from the Aniseed database (Dardaillon et al., 2020) and from the present study, a list of 68 genes with expression in the palps lineage in *Ciona* and/or *Phallusia* (Table S3). We plotted the results of our *Phallusia* RNA-seq data, and found that 70% of the genes were regulated by BMP signaling. Most of them were repressed by BMP, but 20 genes were activated by BMP, and a smaller fraction was repressed or activated depending on the stage. Consequently, the precise function of BMP that is likely to be dynamic in the course of palps differentiation needs to be further investigated in details. Interestingly, Notch is likely to play a role in the specification of the different cell types that compose the palps. For instance, it has been shown that activating Notch represses palps neuronal markers in *H. roretzi* (Akanuma et al., 2002). We found 30 genes regulated by Notch in our dataset.

## Discussion

We have shown that BMP signaling regulates two distinct steps of palps formation in *C. intestinalis*: ANB specification and papilla *vs* inter-papilla specification. Moreover, we have shown conservation of gene expression and regulation by BMP in *P. mammillata*.

### Signaling pathway inhibition and ANB specification

ANB specification is regulated by inputs from several signaling pathways: FGF, Wnt and BMP. While FGF is positively required early on, at the time of neural induction (32-cell stage), all 3 pathways are inactive at the time of ANB fate acquisition as revealed by the expression of *Foxc* (mid-gastrula). This situation is reminiscent of data from vertebrates where anterior neural fate is determined by the triple inhibition of BMP, Nodal and Wnt pathways (Andoniadou and Martinez-Barbera, 2013; Niehrs et al., 2003; Wilson and Houart, 2004). It would thus be interesting to test the function of Nodal inhibition in ANB specification since we have already shown that it is involved in posterior neural fate determination in *Ciona* (Roure et al., 2014). While it appears that active FGF, Wnt or BMP signaling is incompatible with ANB determination, the specific function of each pathway seems different. FGF appears to regulate anterior CNS *vs* ANB fate decision along the antero-posterior axis (Wagner and Levine, 2012). Wnt seems to regulate *Foxc*+ ANB fate *vs Foxc*-ANB fate along the medio-lateral/dorso-ventral axis (Feinberg et al., 2019). Finally, BMP might participate in the segregation between ANB and immediately anterior/ventral epidermal fates. Finer details on the function of these pathway in ANB fate determination and on their likely cross-talk should be an exciting line of research in this simple and geometric model system.

### From ANB to palps differentiation

Our results of late inhibition of BMP signaling (from gastrula stages) indicate that *Foxg*, expressed in a single row of cells with a U-shape, delineates cells competent to become papilla. A network of gene interactions has previously been identified that regulates the transition of *Foxg* from a U-shape to 3-spots eventually forming protruding papillae (Liu and Satou, 2019). BMP is an input to this network, presumably through the regulation of *Sp6/7/8/9* expression. Importantly, we have shown that BMP signaling is active in the median palps forming region before the onset of *Foxg* and *Sp6/7/8/9* expression. This region most likely corresponds to the future ventral palp. However, activating BMP at this stage, does not result in ectopic palp formation but to an absence of the ventral palp. This discrepancy might be better understood by more finely controlling levels of BMP signaling but also its timing and cells that receive it, through optogenetics for example. Nevertheless, our observations point to differences between the two symmetrically bilateral dorsal palps and the single ventral and median palp. While we are not aware of ventral palp-specific marker, we have shown that *Msx* is transiently expressed only in the ventral palp (Fig 6); this may also be the case for *Hes*.*a* (Chowdhury et al., 2022). In addition, the *Pou4*+ sensory neurons are located dorsally in the ventral palp, while they are located around *Isl*+ cells in the dorsal palps. Specific dorsal and ventral genetic sub-network would thus be interesting to uncover.

### Conservation of PNS formation in chordates?

The ascidian larval PNS, palps included, originates from the neural plate border with the exception of the ventral tail PNS that originates from a region at the opposite end of the embryo, the ventral epidermis. Signaling pathways are pleiotropic and are consequently poor indicators of possible evolutionary conservation. Nevertheless, it is striking that FGF, Wnt and BMP are deployed in ascidians to regulate neural plate border specification and differentiation of its derivatives, most likely with changing dynamic requirements at diverse developmental stages. This is reminiscent of the mechanisms regulating neural plate border and its derivatives, the cranial placodes and the neural crest (Martik and Bronner, 2021; Pla and Monsoro-Burq, 2018; Stundl et al., 2021). The similarities extend beyond signaling pathways since a suite of genes have conserved expression between ascidians and vertebrates, and have led to several evolutionary scenarios (Cao et al., 2019; Horie et al., 2018; Pasini et al., 2006; Poncelet and Shimeld, 2020). Our present study add material to gene network level comparisons.

The ascidian PNS is made of epidermal sensory neurons that have different morphologies, connectivity and sensory capacities depending on their location (Abitua et al., 2015; Imai and Meinertzhagen, 2007; Ryan et al., 2018). However, they share a number of genes marking the presumptive domains or differentiating neurons. Yet, what regulates their specific identities is still incompletely understood (Chacha et al., 2022). For example, a number of genes expressed in the tail PNS are also expressed in the palps, and these expression domains are conserved in species that have diverged almost 400 My ago (Table S3) (Akanuma et al., 2002; Coulcher et al., 2020; Joyce Tang et al., 2013; Pasini et al., 2006; Roure and Darras, 2016). Comparative approaches of PNS formation between divergent ascidian species and across chordates (vertebrates and cephalochordates) promise to yield insights into PNS evolution and the flexibility of developmental mechanisms.

## Materials and methods

### Embryo obtention and manipulation

Adults from *Ciona intestinalis* (formerly referred to *Ciona intestinalis* type B (Brunetti et al., 2015)) were provided by the Centre de Ressources Biologiques Marines in Roscoff (EMBRC-France). Adults of *Phallusia mammillata* were provided by the Centre de Ressources Biologiques Marines in Banyuls-sur-mer (EMBRC-France) following diving or by professional fishermen following trawling in the Banyuls-sur-mer (France) area. Gametes collection, *in vitro* fertilization, dechorionation and electroporation were performed as previously described (Coulcher et al., 2020; Darras, 2021); and staging of embryos was performed according to the developmental table of *Ciona robusta* (Hotta et al., 2007).

Electroporation constructs used in this study have been previously described (Pasini et al., 2006). Embryos were treated with 150 ng/ml of recombinant mouse BMP2 protein (355-BEC, R&D Systems Inc, 100 μg/mL stock solution in HCl 4 mM + BSA 0.1 %), 100 ng/ml of recombinant human bFGF (F0291, Sigma-Aldrich, 50 μg/mL stock solution in 20 mM Tris pH=7.5 + BSA 0.1 %) complemented with 0.1% BSA, or 2.5 μM of the BMP receptor inhibitor DMH1 (S7146, Euromedex, 10 mM stock solution in DMSO) at the stages indicated in the text and figures. Control embryos were incubated with sea water containing 0.1 % BSA and/or 0.025% DMSO.

### *In situ* hybridization and immunostaining

For all labeling experiments, embryos were fixed in 0.5 M NaCl, 100 mM MOPS pH=7.5 and 3.7% formaldehyde. Whole mount chromogenic *in situ* hybridization were performed using plasmid cDNA or synthetic DNA (eBlocks Gene Fragment, IDT) as templates for probe synthesis (Tables S2 and S4) as described previously (Chowdhury et al., 2022). Gene models and identifiers correspond to the following genome assemblies, KH2012 for *Ciona robusta* (Satou et al., 2008) and MTP2014 for *Phallusia mammillata*, that were retrieved from the Aniseed database (Dardaillon et al., 2020). Images were acquired using an AxioCam ERc5s digital camera mounted on a stereomicroscope (Discovery V20, Zeiss). The number of experiments and embryos for phenotypic effects by gene expression analysis are shown in the figures and their legends.

Fluorescent *in situ* hybridization were adapted from (Racioppi et al., 2014). Briefly, digoxigenin-labeled probes were recognized using an anti-DIG antibody coupled to peroxidase (11207733910, Roche), and fluorescein-labeled probes were recognized using an anti-FLUO antibody coupled to peroxidase (11426346910, Roche). Fluorescence signal was produced using the TSA plus kit (NEL753001KT, Perkin-Elmer) following manufacturer’s recommendations with cyanin3 and fluorescein for DIG- and FLUO-probes respectively.

Active BMP signaling was visualized by immunostaining using a rabbit monoclonal antibody against mammal Smad1, Smad5 and Smad8 phosphorylated at two serine residues at the C-terminal end (clone 41D10, #9516, Cell Signaling Technology) diluted at 1:200. The epitope is present in the single ortholog Smad1/5/8 of both *Ciona intestinalis* and *Phallusia mammillata*. Anti-rabbit coupled to Alexa Fluor 568 (A11011, Invitrogen) was used at 1:400 for visualization. Similar data were obtained using another antibody (clone D5B10, #13820, Cell Signaling Technology) (data not shown). Membranes were stained using Alexa Fluor 594 phalloidin (A12381, Invitrogen) used at 1:1000. Nuclei were stained using DAPI. Image acquisition was performed using confocal microscopy (Leica SP8-X, BioPiC platform, Banyuls-sur-mer). Confocal z-stacks were visualized and analyzed in 3D using the Imaris 8.3 software (Bitplane). In particular, this software was used to count the number of cells expressing a gene of interest. In brief, fluorescent signals were converted as 3D objects: *in situ* hybridization signals as surface objects, and DAPI-labeled nuclei as spots. The number of spots within a given surface was used as a proxy for the number of cells expressing a gene. Snapshots of such analyses and 3D renderings are shown in Fig 1, 3 and 6. Image panels and figures were constructed with Affinity Photo and Affinity Designer.

## Acknowledgements

We thank G. Diaz (Port-Vendres) and staff (divers and boat crew) at the marine stations of Banyuls-sur-mer and Roscoff (French node of the European research infrastructure EMBRC) for providing animals. We would like to acknowledge the BioPiC imaging facility (Sorbonne Université/CNRS, Banyuls-sur-mer), R. Dumollard and H. Yasuo for sharing plasmids, and V. Thomé for advices on fluorescent *in situ* hybridization and immunostaining.

## Authors’ contributions

AR and SD designed the project. AR performed most of the experiments and analyses with the help of RC and SD. SD supervised the project, wrote the manuscript and obtained funding. All authors edited the manuscript, read and approved the final version.

## Data availability

RNA-seq data are available under the BioProject ID PRJNA779382. All other data generated or analyzed during this study are included in the manuscript and supporting files.

## Supplementary data

**Figure S1.**
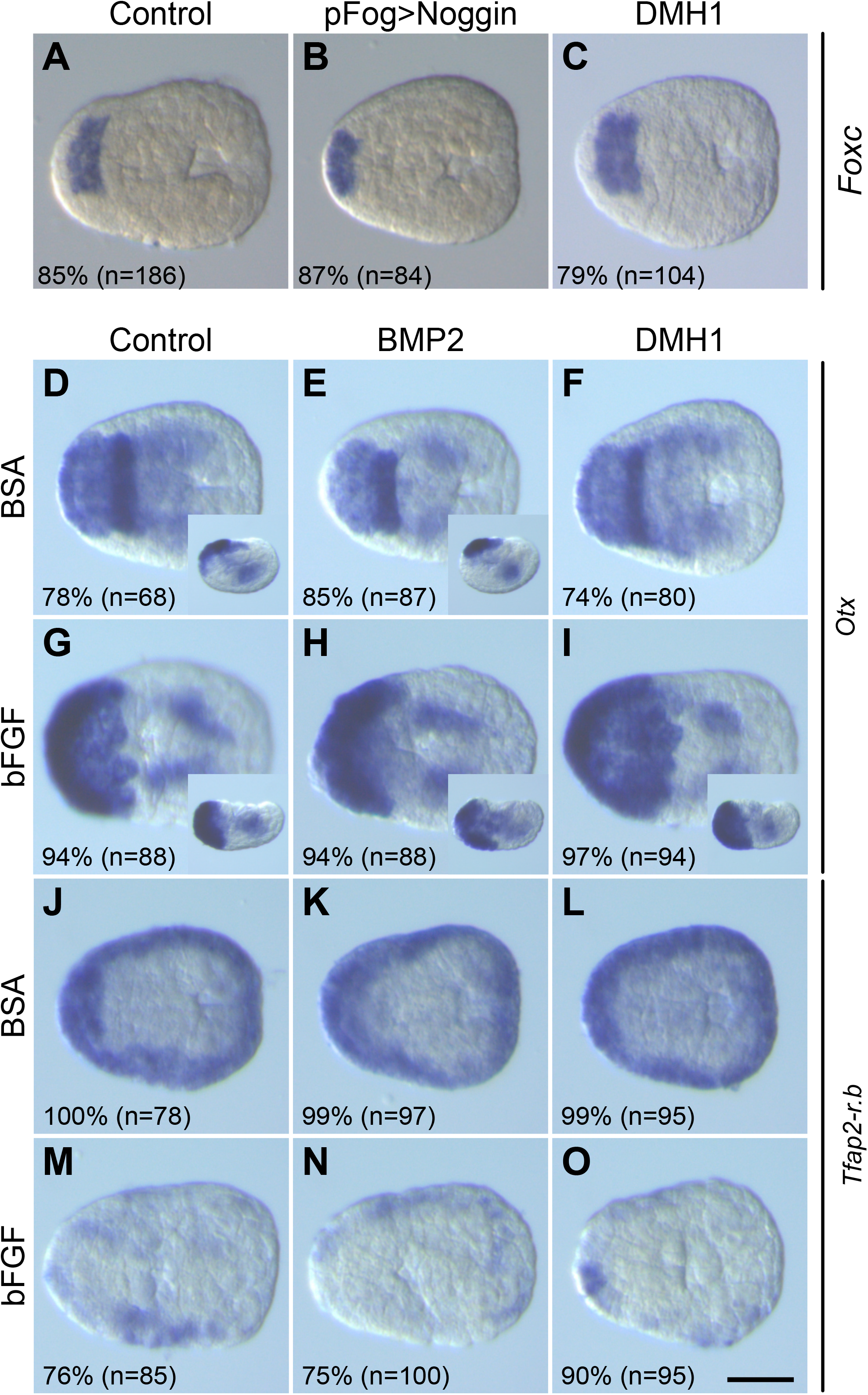
Expression of neural and epidermal markers following BMP and FGF pathways modulations. (A-C) *Foxc* expression by *in situ* hybridization at early neurula stages (St. 14) is unchanged following BMP pathway inhibition by Noggin overexpression (B) or DMH1 treatment from the 8-cell stage (C). (D-O) Embryos were treated from the 8-cell stage to the fixation at early neurula stages (St. 14) with 100 ng/ml recombinant BMP2 protein or 2.5 μM DMH1 alone, or in combination with 100 ng/ml recombinant bFGF protein. Gene expression was assessed by *in situ* hybridization for *Otx* (D-I) and *Tfap2-r*.*b* (J-O). For each panel, n indicates the number of embryos examined. The percentages indicate the frequency of the phenotype depicted in the picture. The results come from two independent experiments or more. Embryos are shown in neural plate views with anterior to the left except insets that are lateral views with anterior to the left and dorsal to the top. Scale bar: 50 μm.

**Figure S2.**
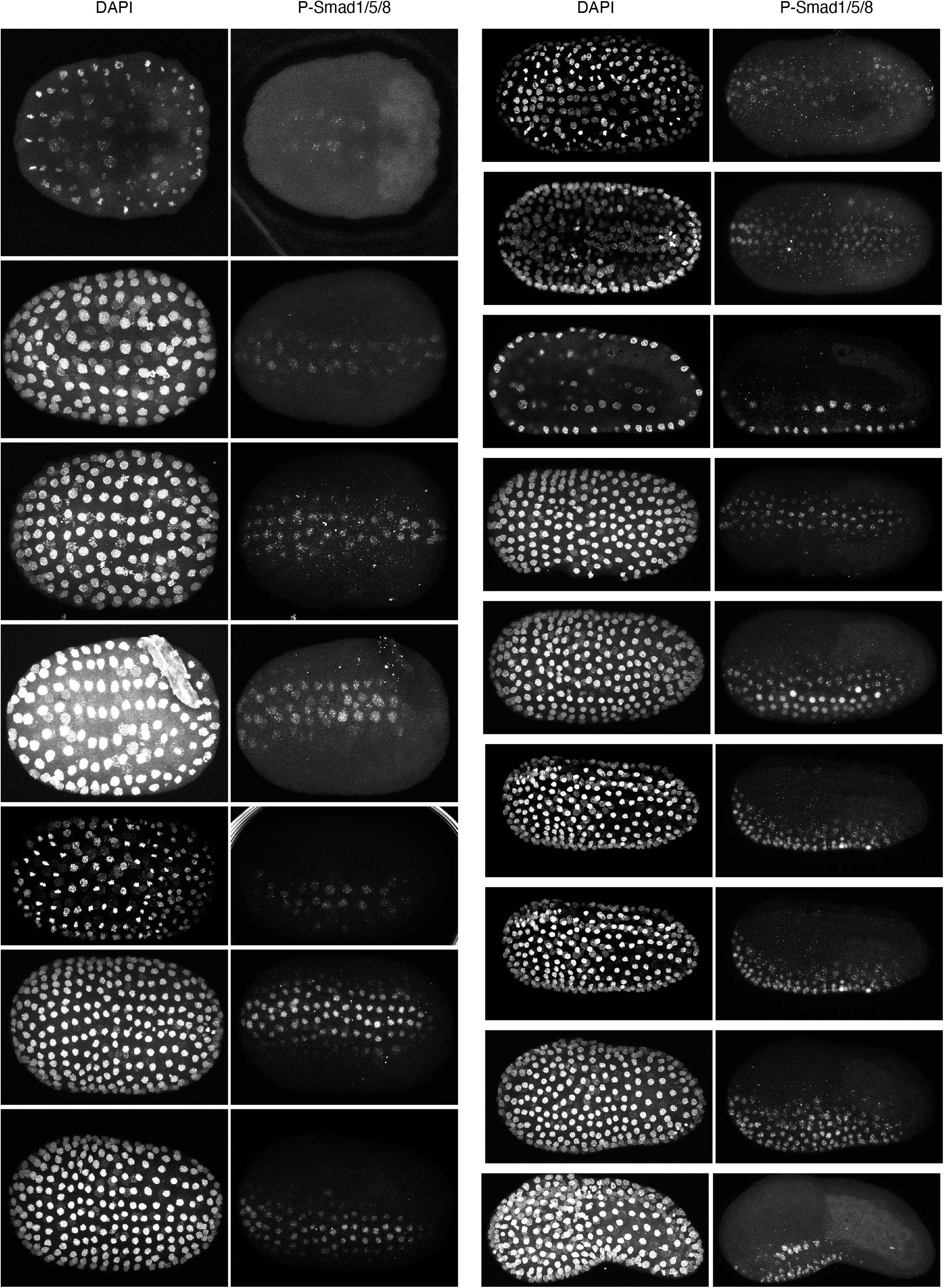
P-Smad1/5 immunostaining in *P. mammillata*.

**Table S1.**
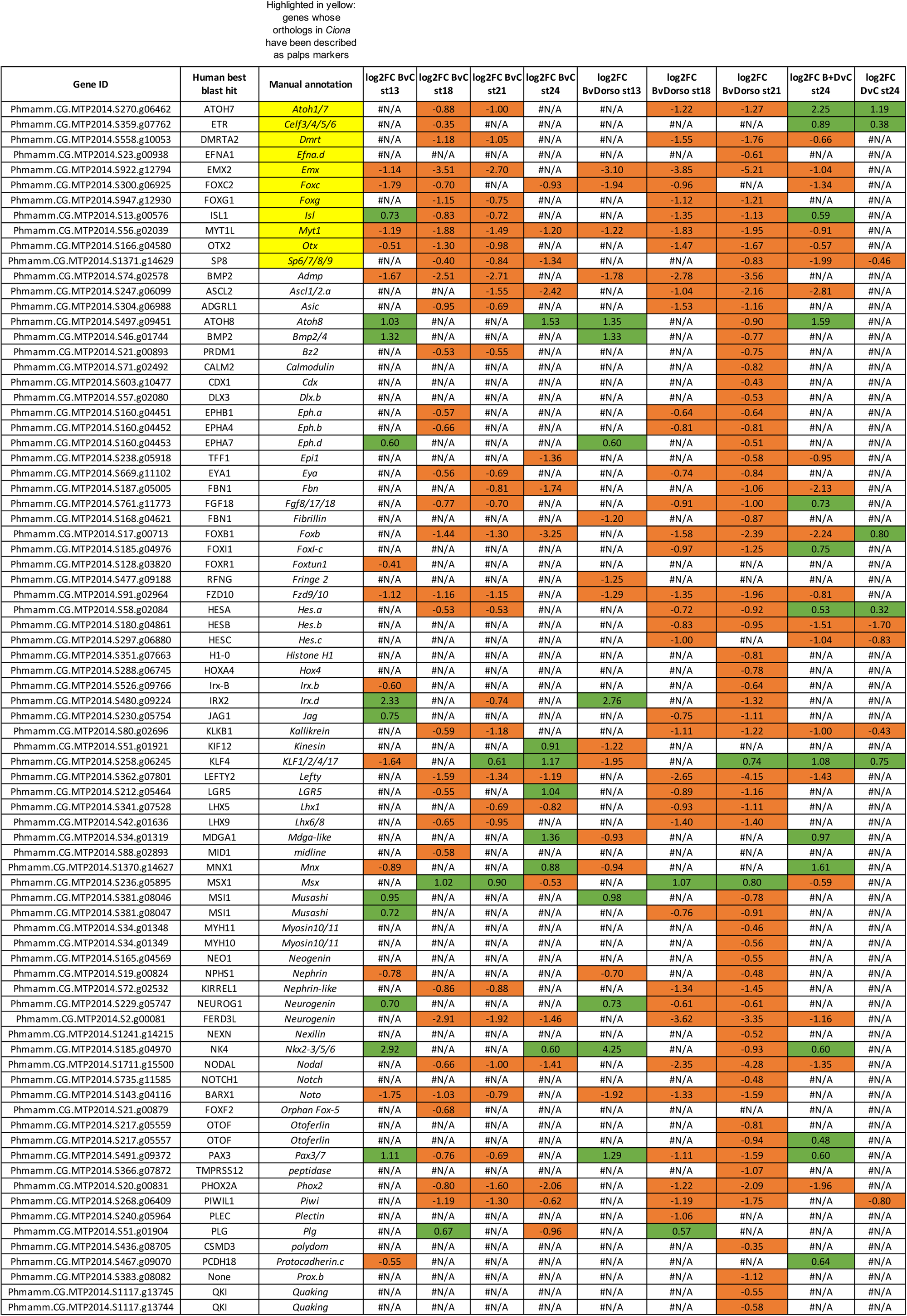
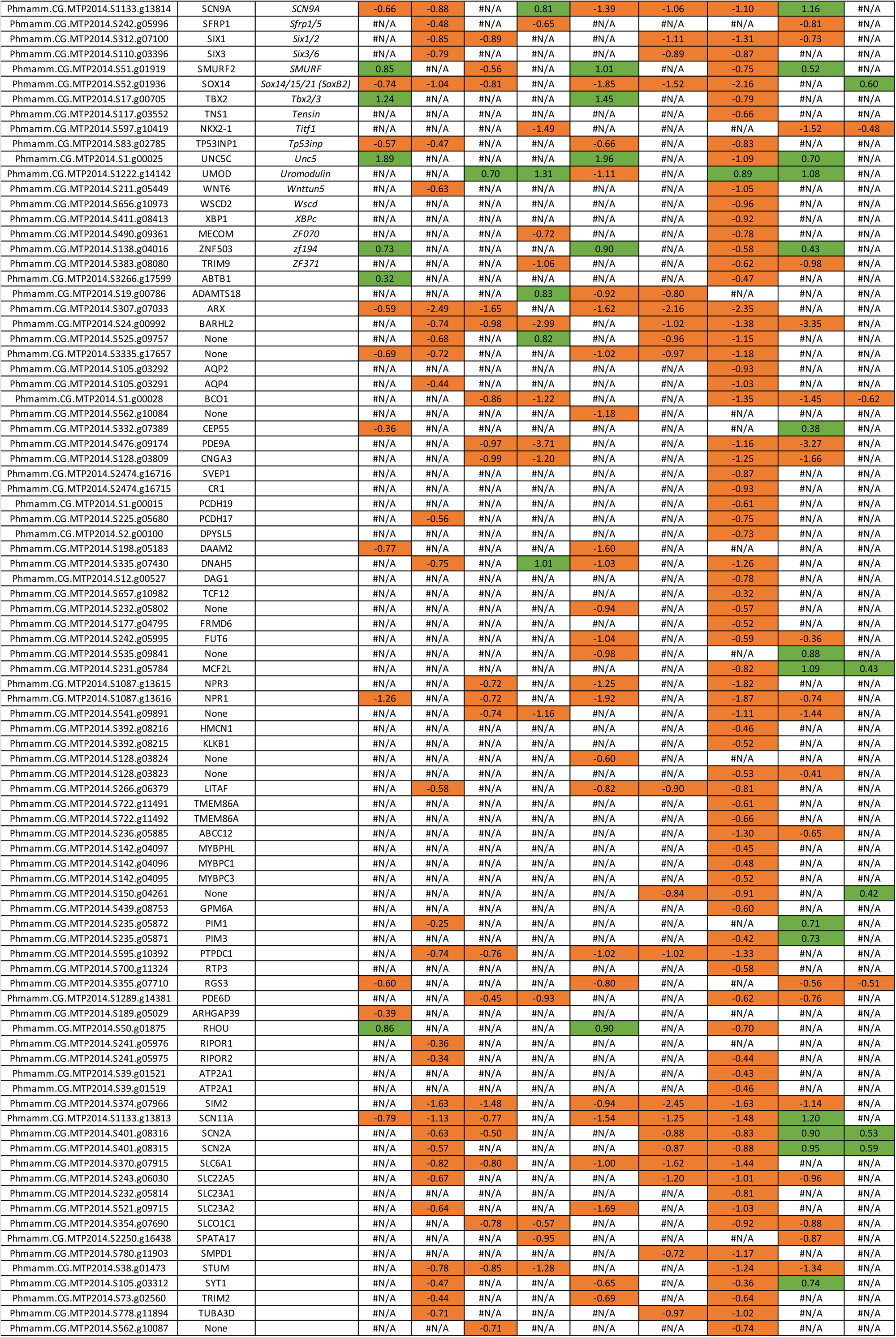
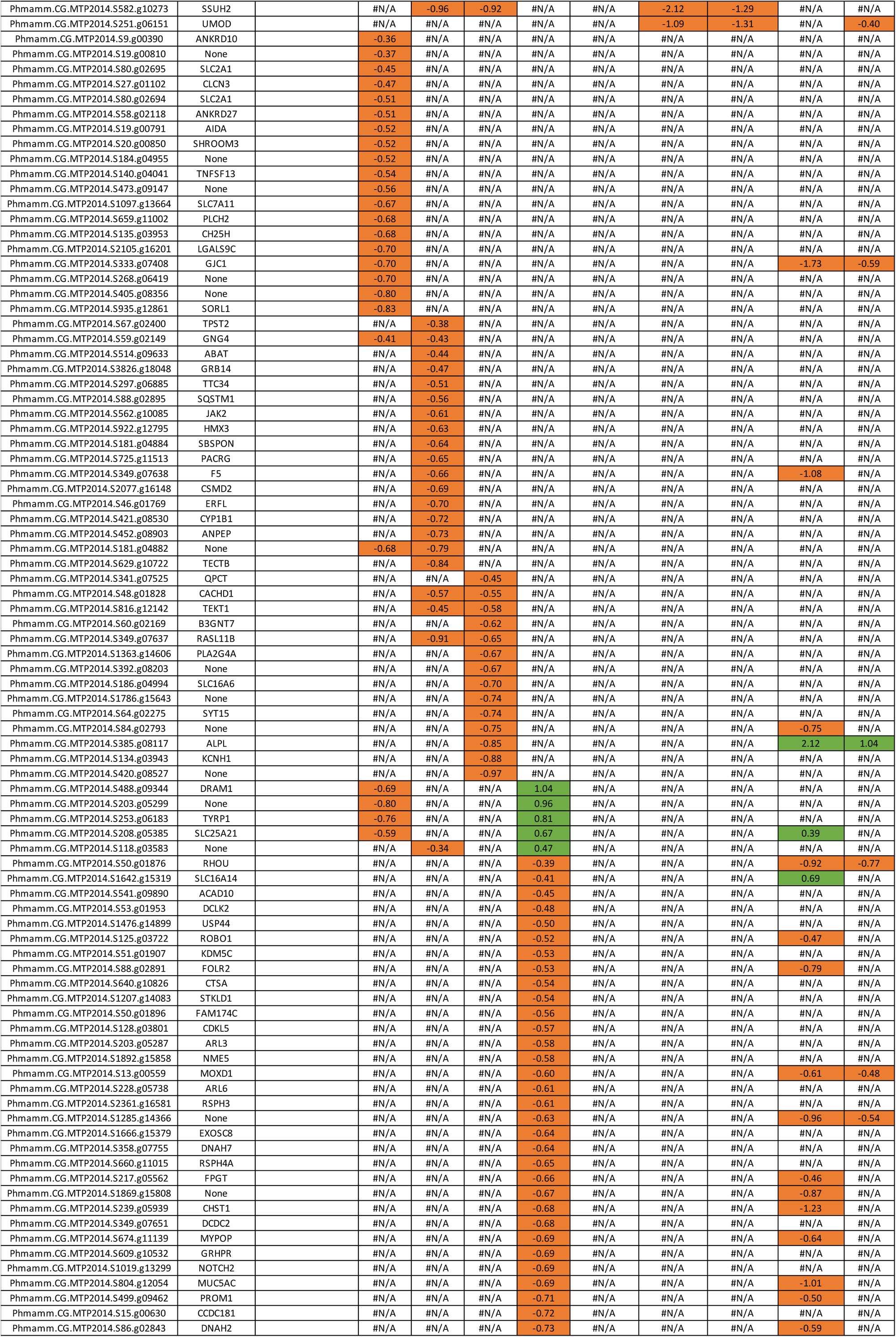
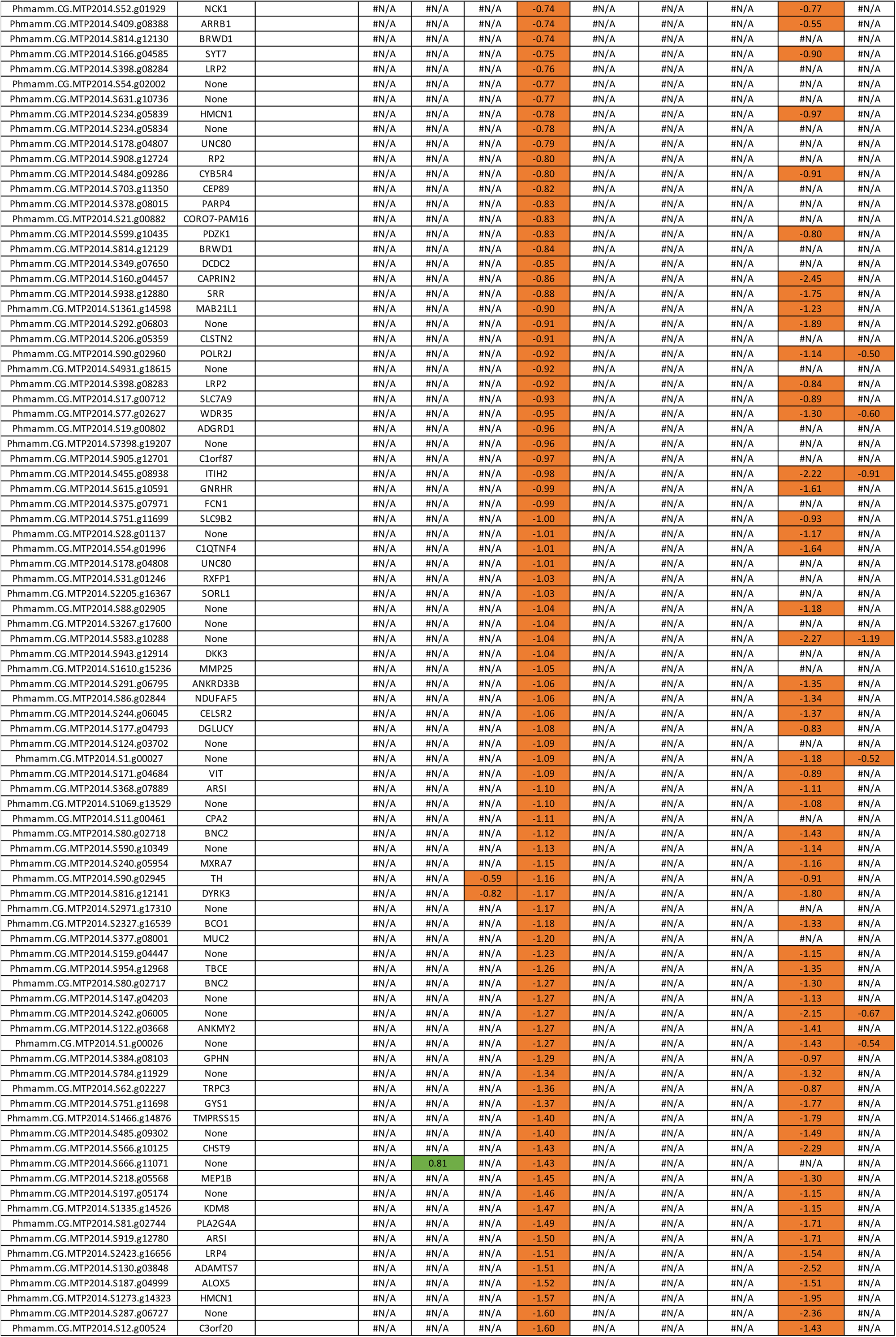
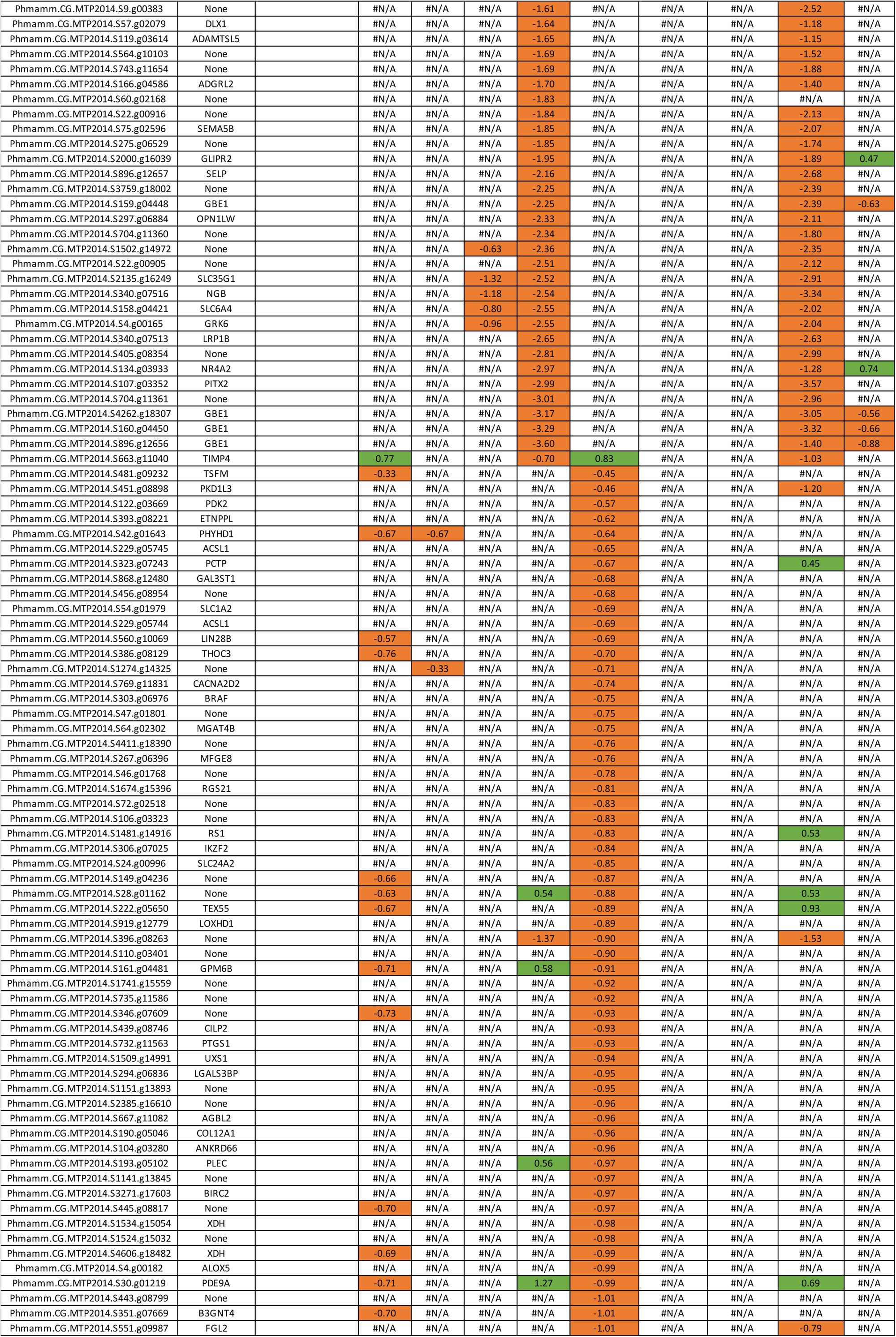
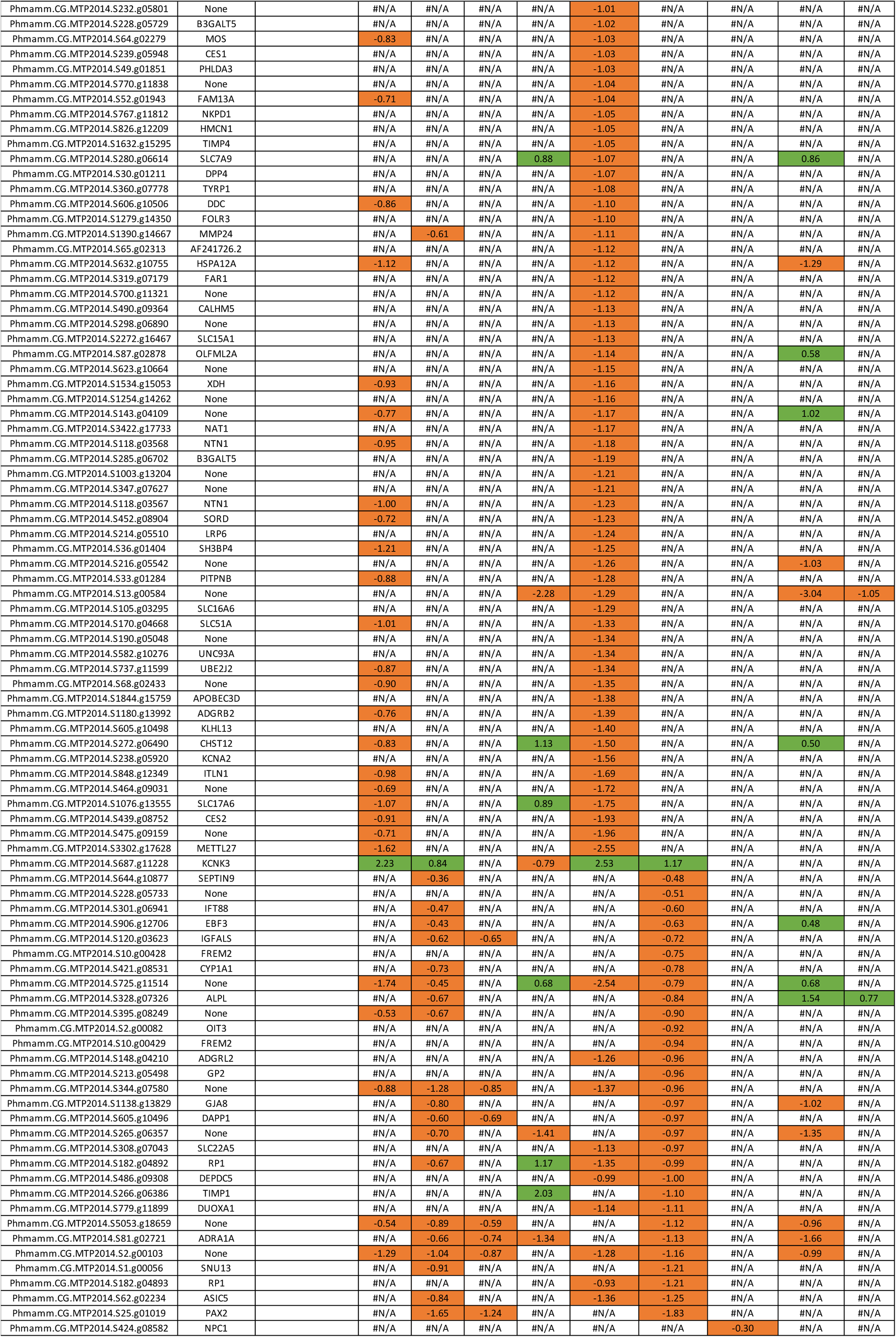
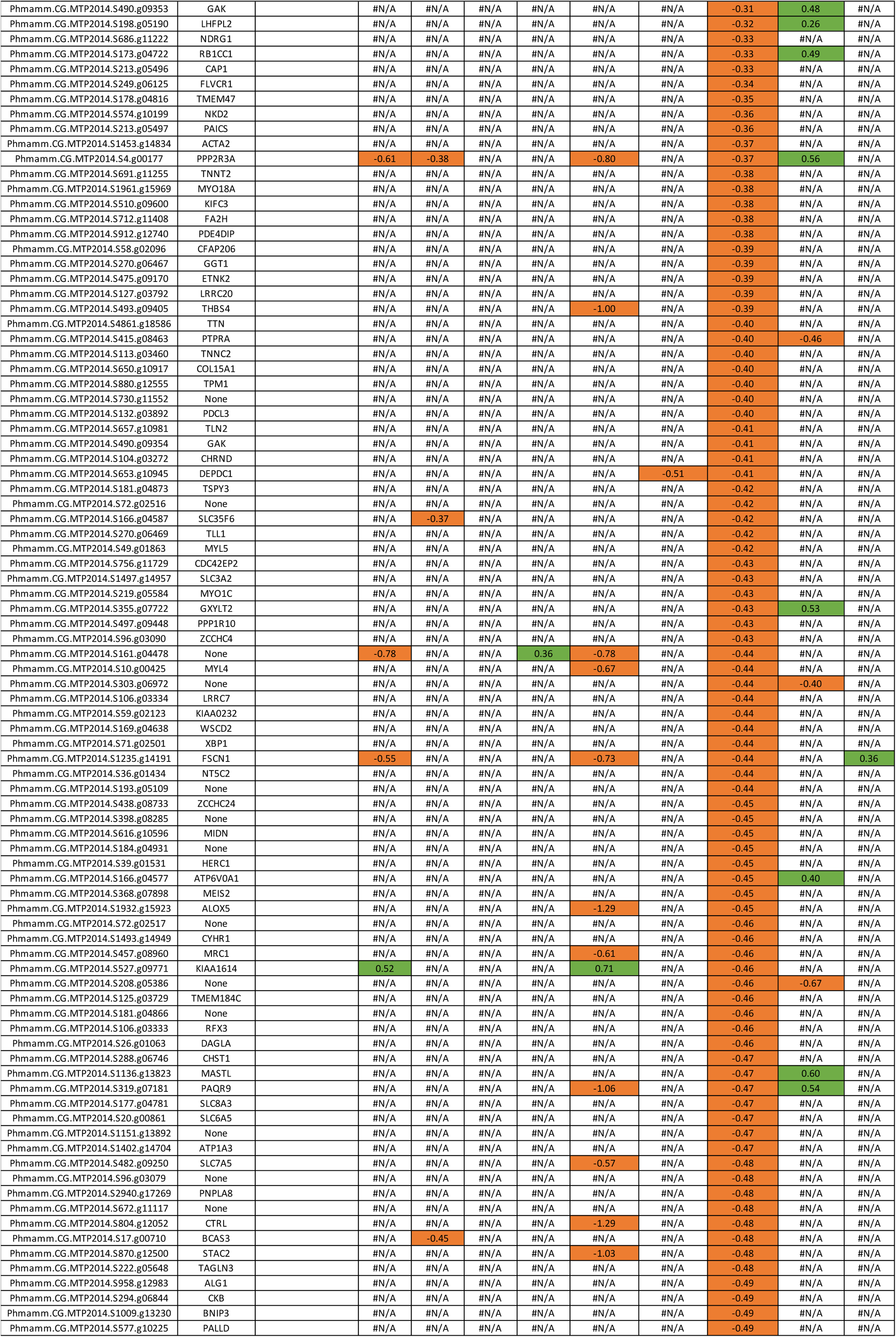
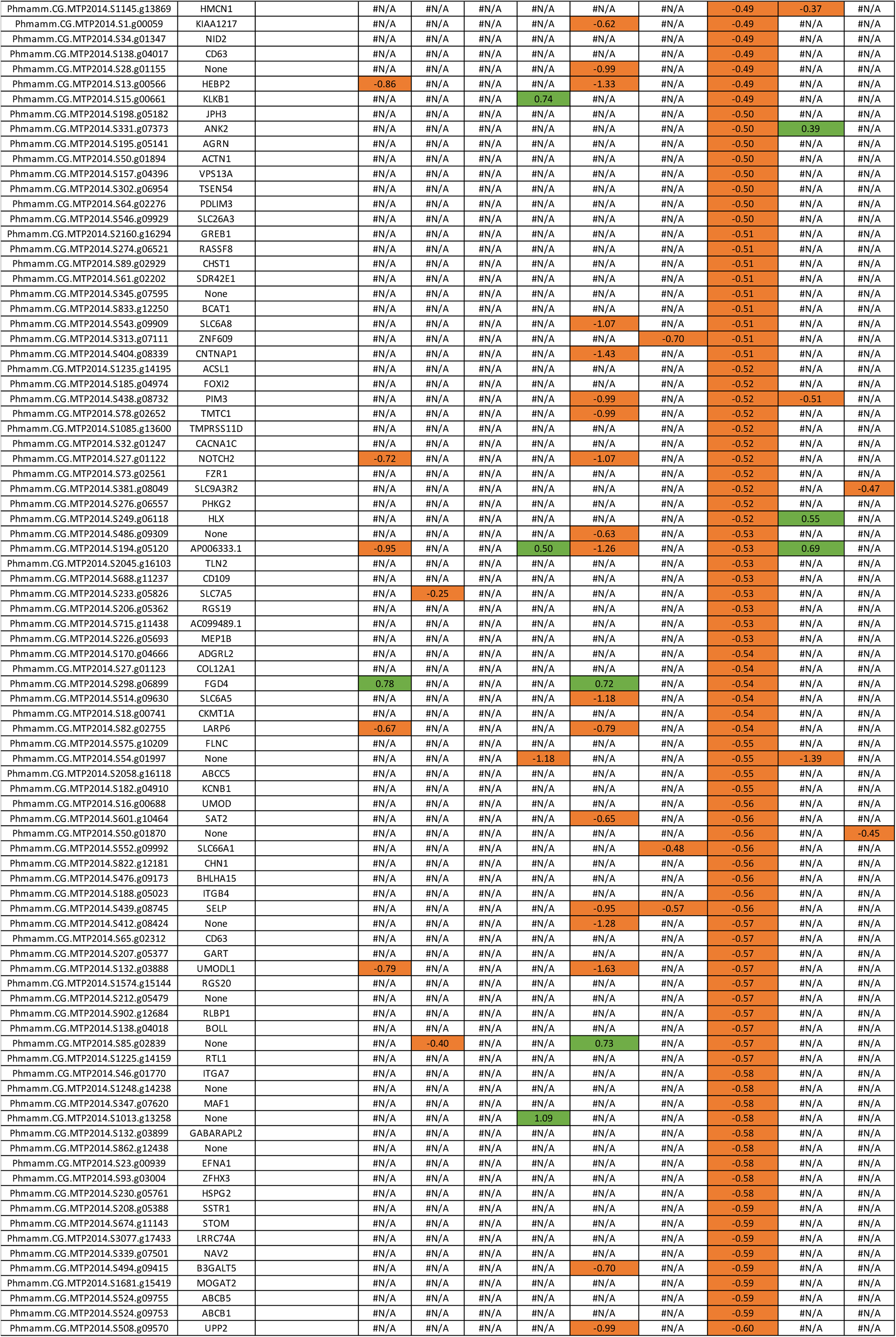
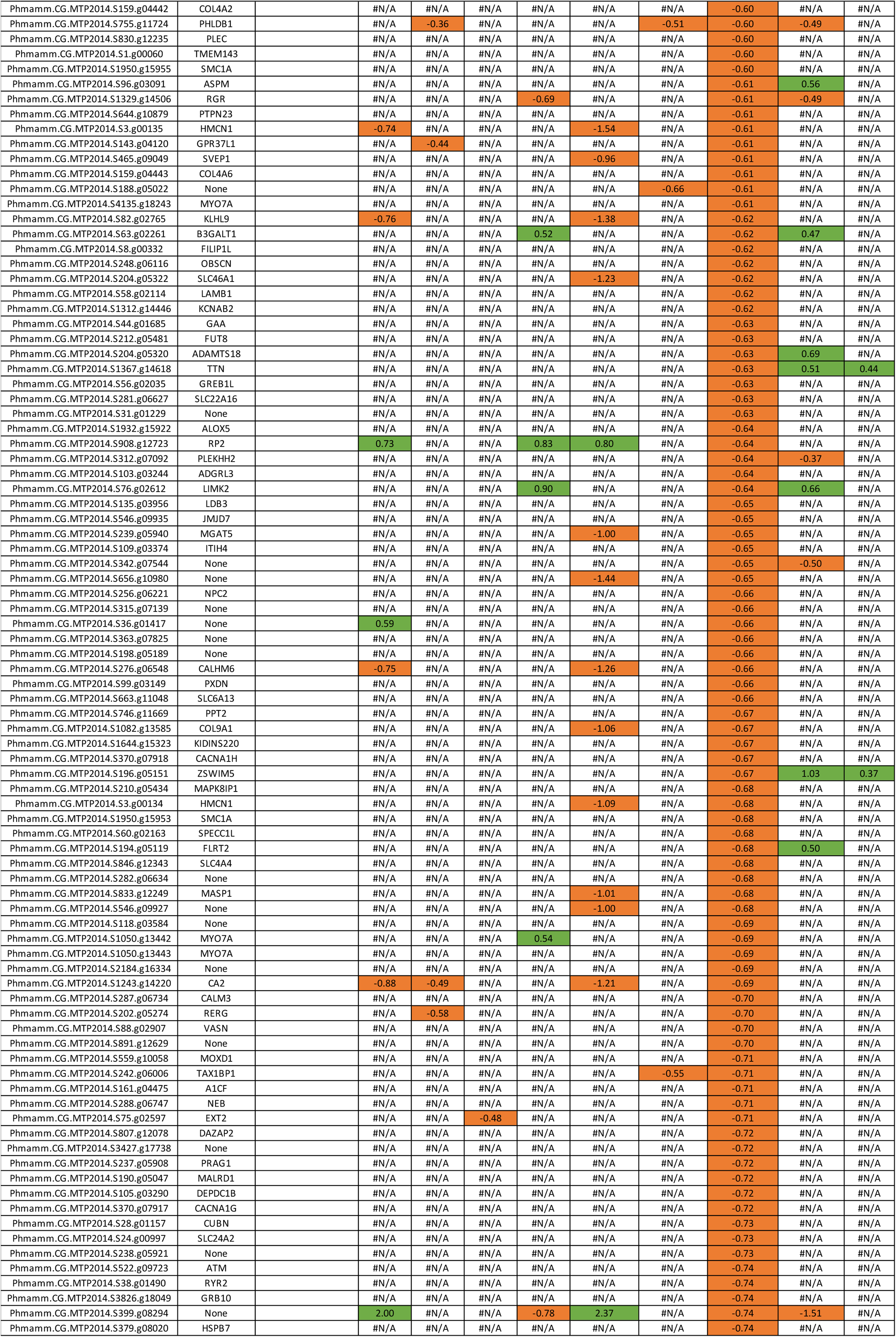
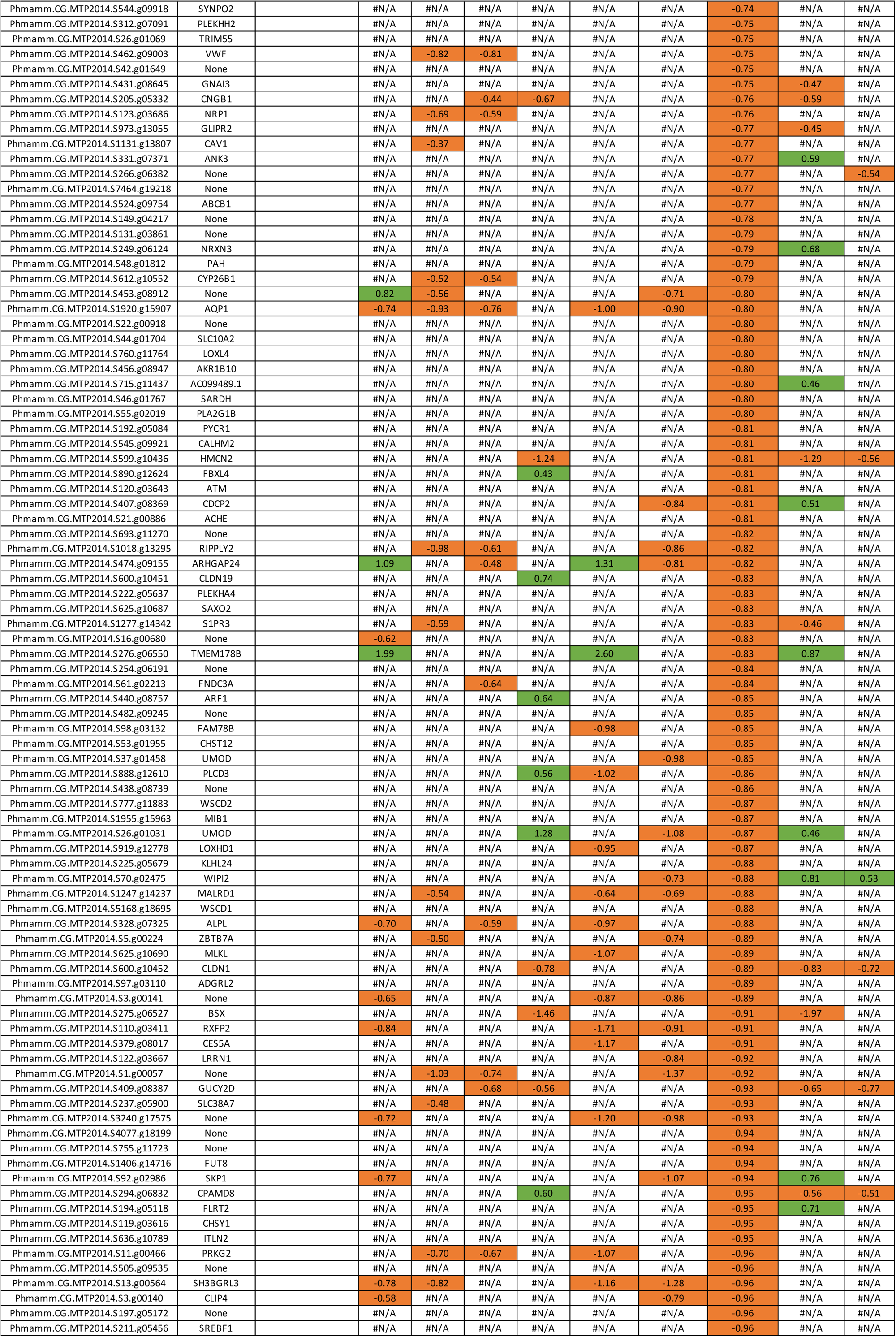
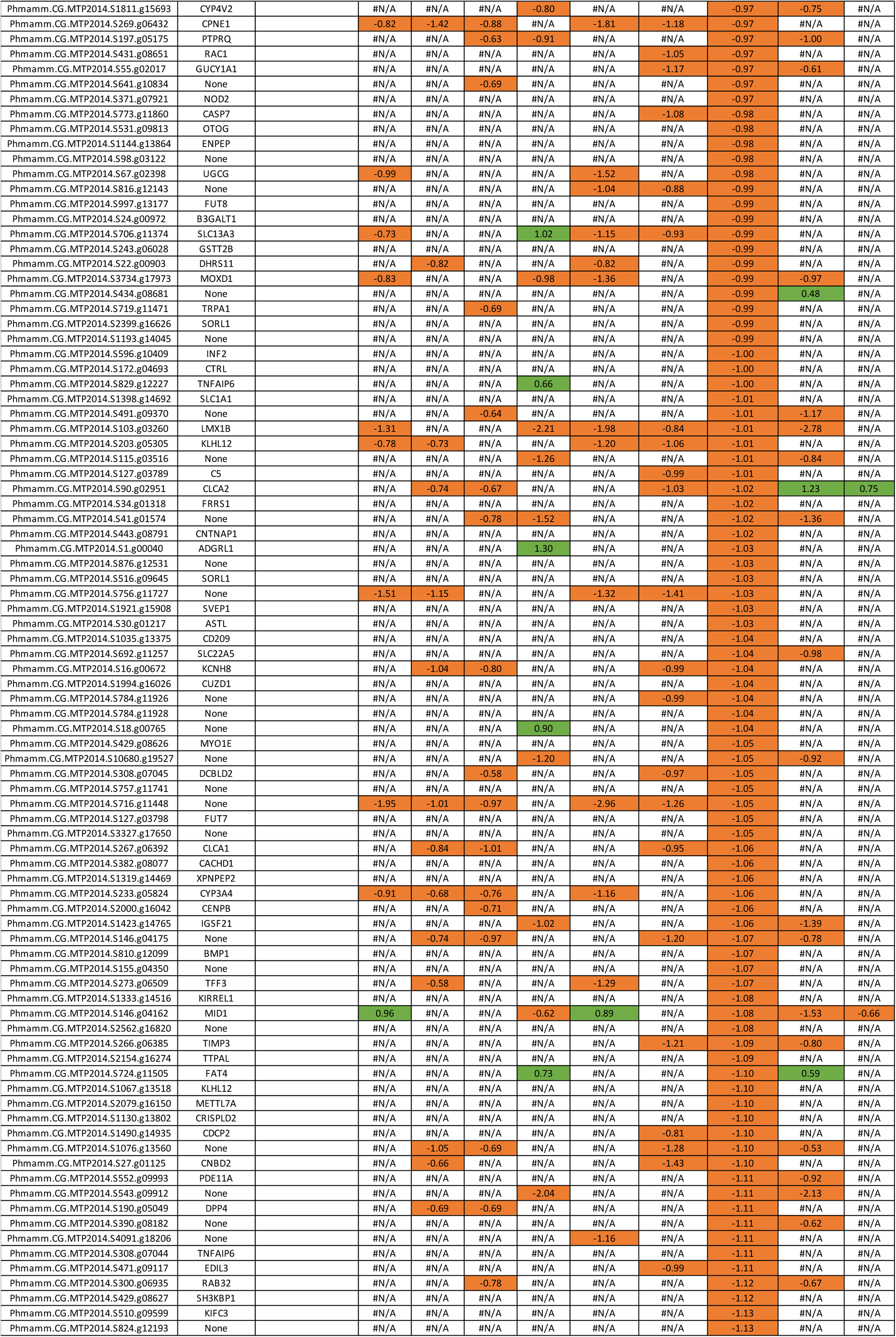
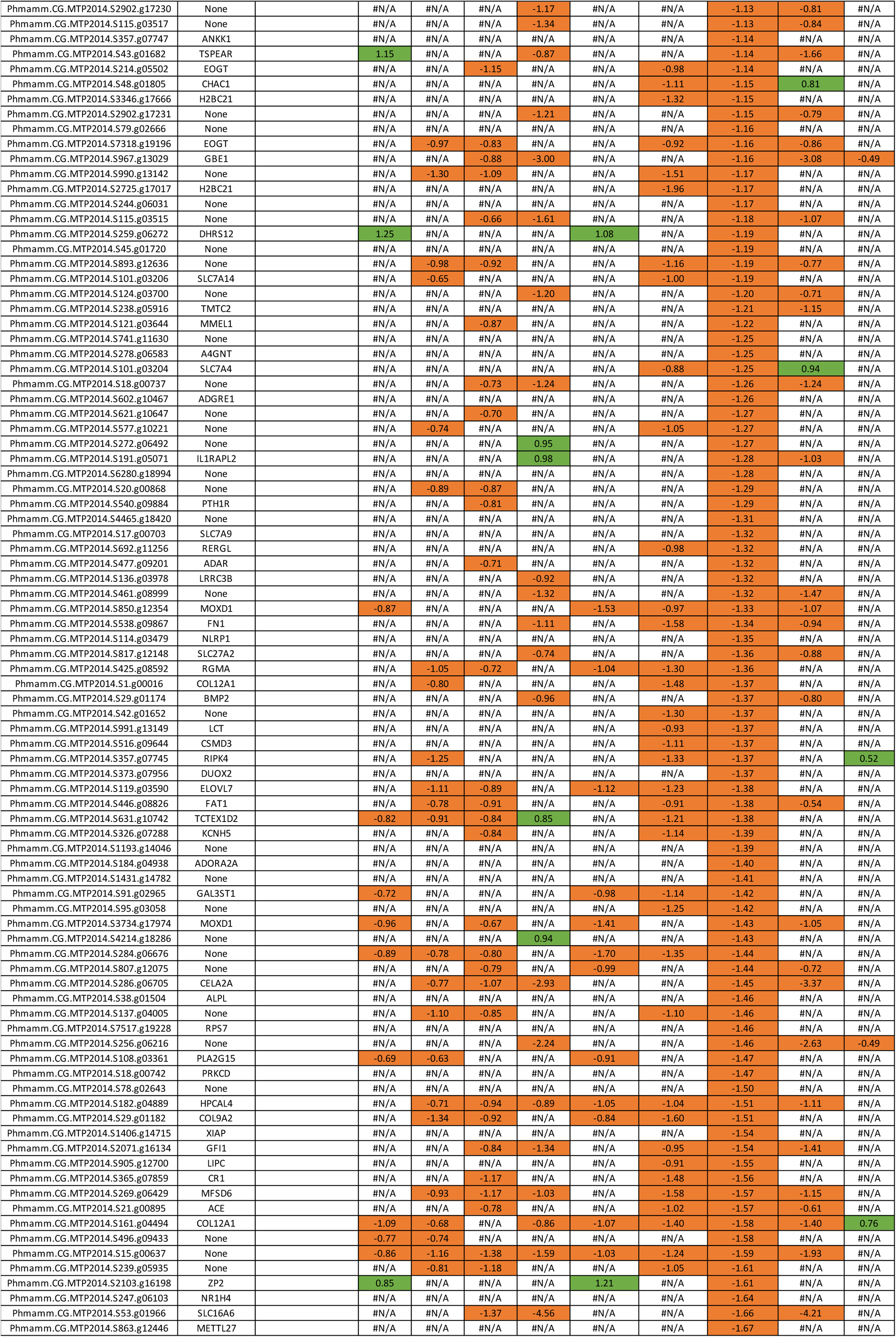
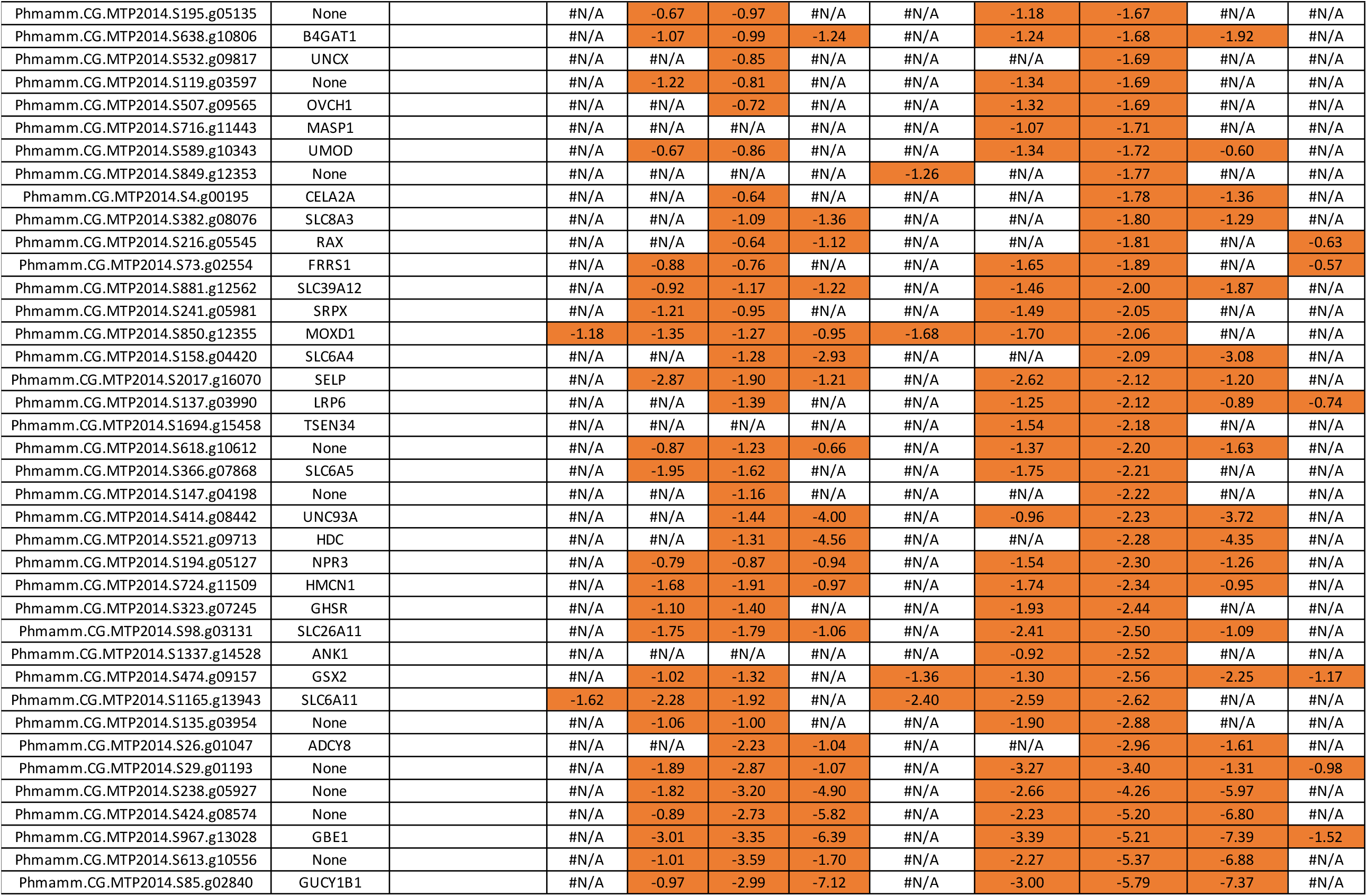
List of the 1098 genes repressed by BMP signaling in *P. mammillata*. Processing of the *P. mammillata* RNA-seq data from the BioProject PRJNA779382 has been described in (Chowdhury et al., 2022). Here are shown the genes with a negative log2 fold change as calculated by DESeq2 with a p-value<0.05. BvC: BMP4 treatment *vs* control. BvDorso: BMP4 treatment *vs* Dorsomorphin treatment. B+DvC: BMP4+DAPT treatment *vs* control. DvC: DAPT treatment *vs* control.

**Table S2.**
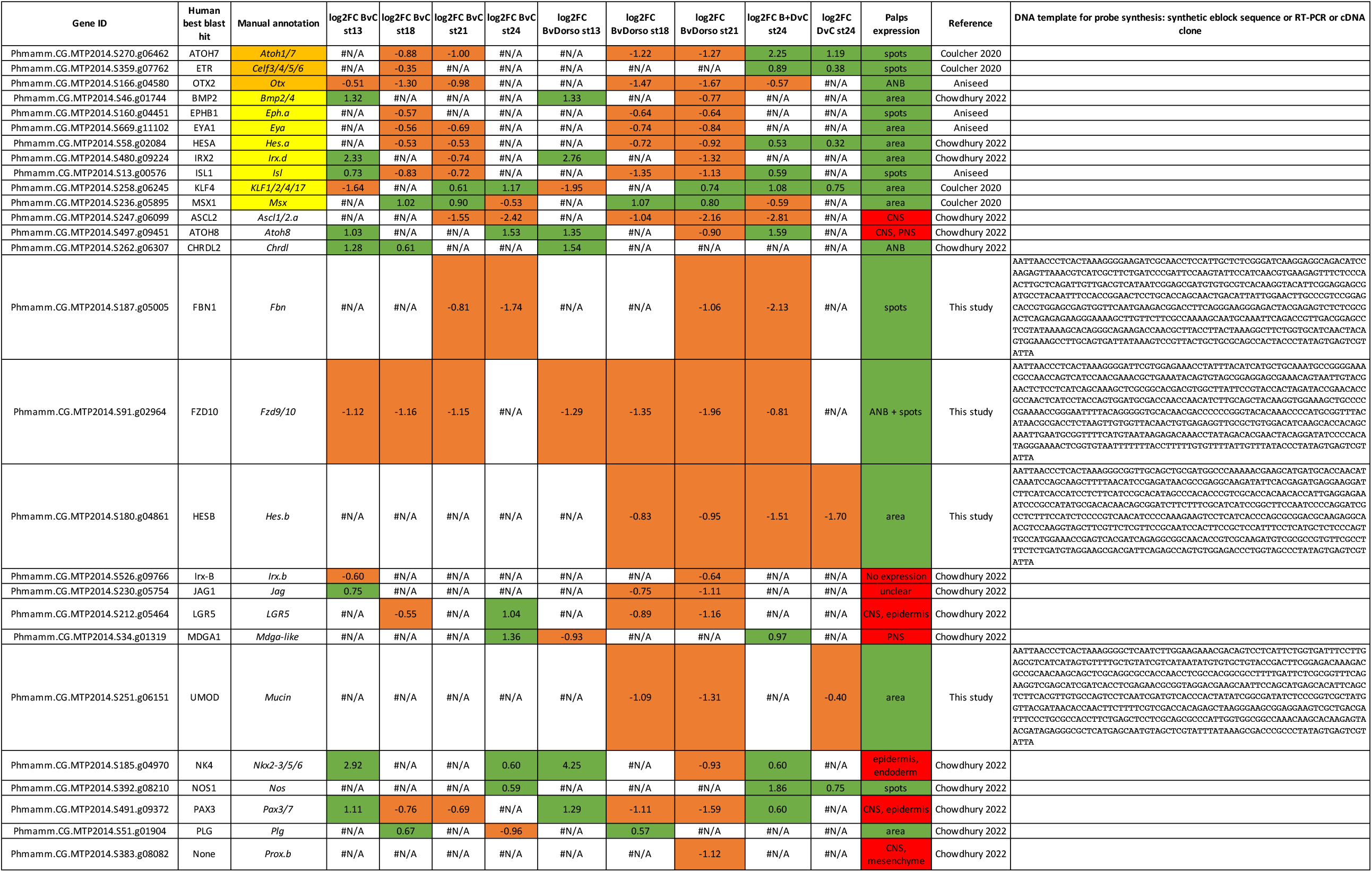
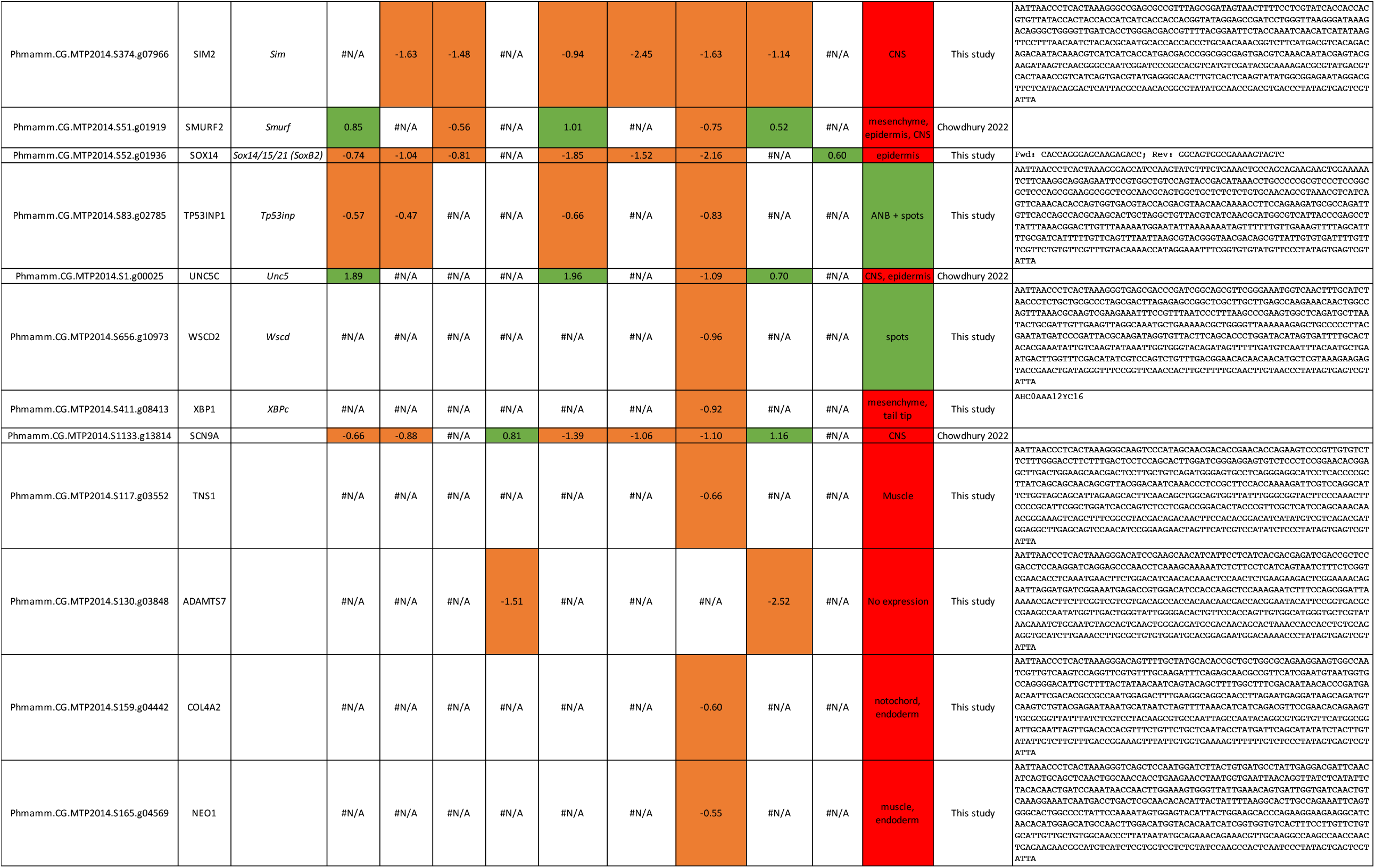
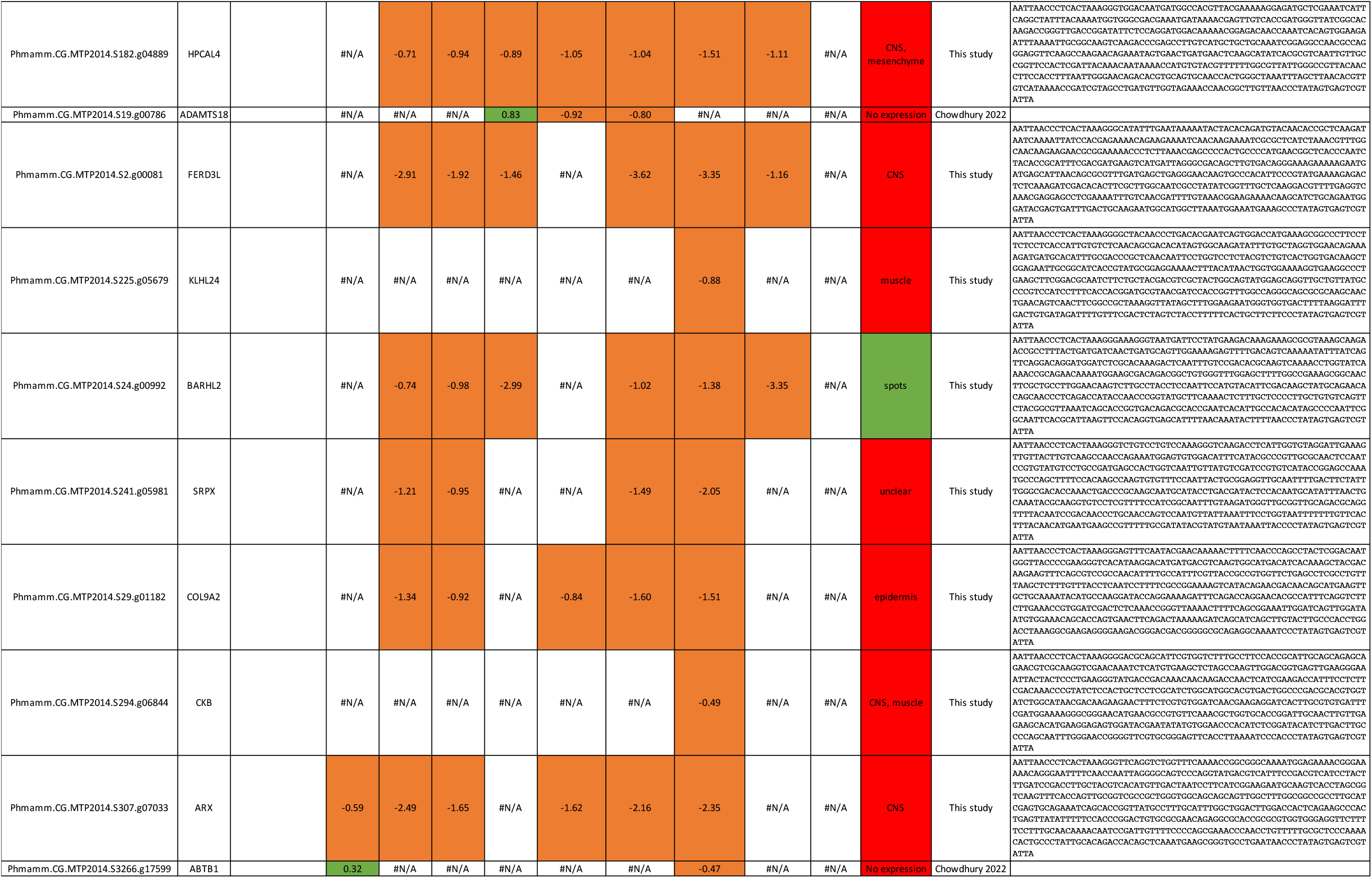
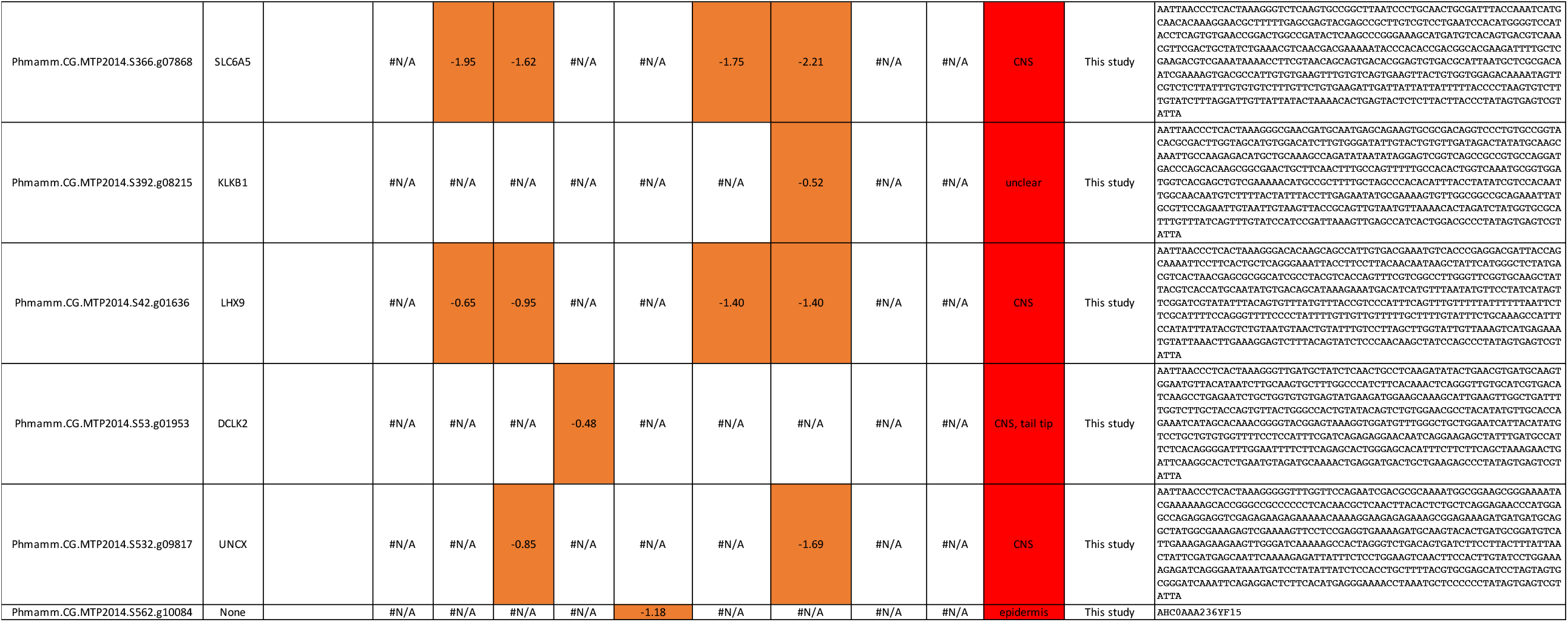
Expression patterns for a list of the 55 genes regulated by BMP signaling in *P. mammillata*.

**Table S3.**
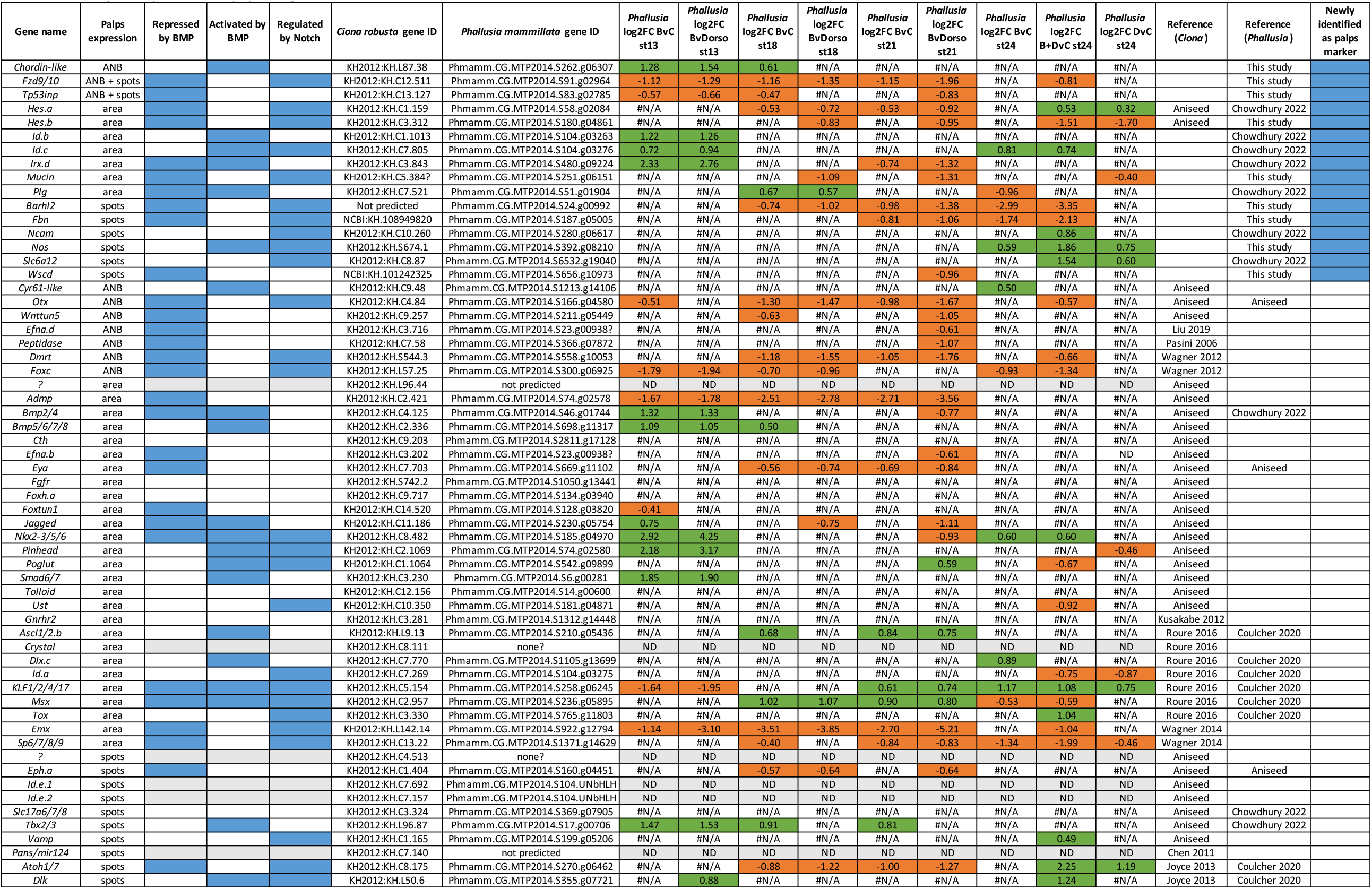
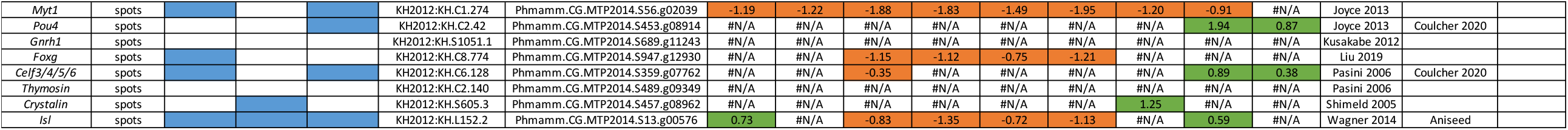
List of 68 genes expressed in the palps lineage in *Ciona* or/and *Phallusia*.

**Table S4.**
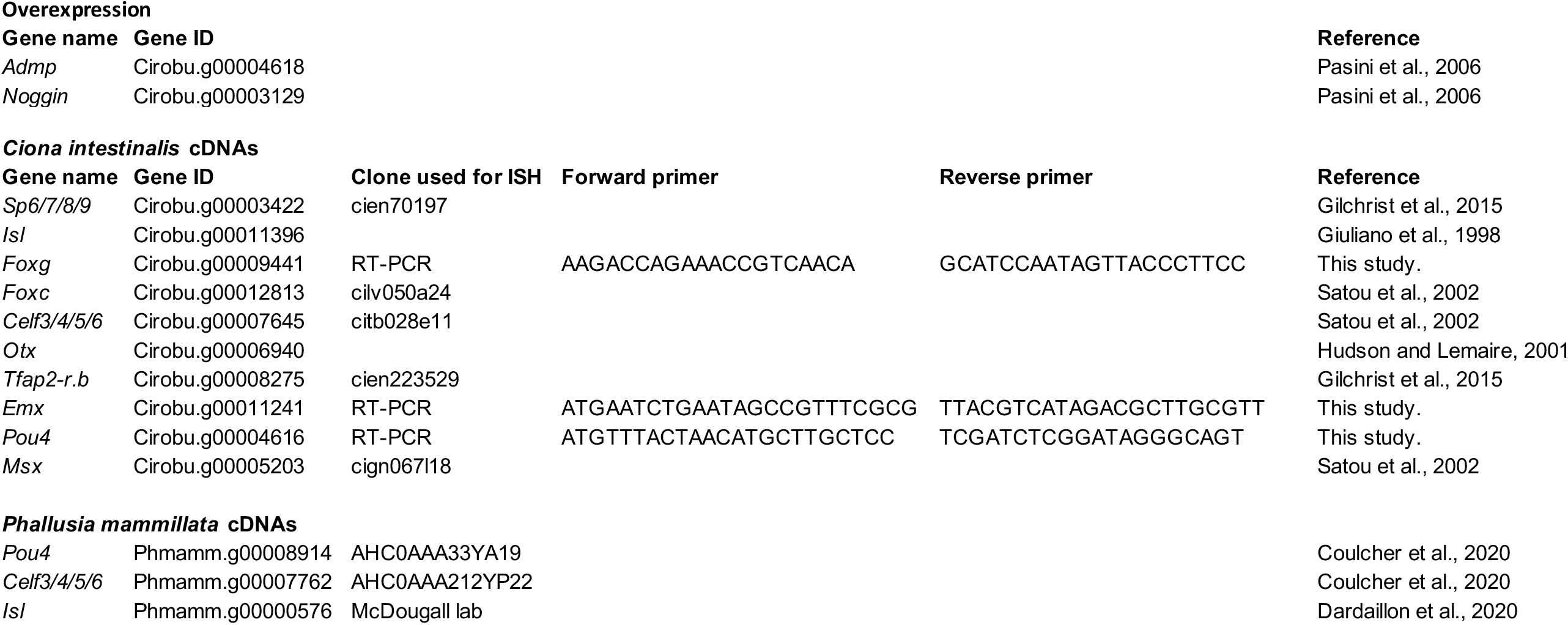
Gene identifiers and references. Unique gene identifiers have been retrieved from Aniseed.

